# Structural basis for the function of long noncoding RNA *Pnky* in neural stem cells

**DOI:** 10.1101/2025.09.01.671568

**Authors:** Parna Saha, Shivali Patel, Rafael CA. Tavares, Hyeonseok Choi, Rebecca E. Andersen, Lucille H Tsao, Anna Marie Pyle, Daniel A Lim

## Abstract

LncRNA *Pnky* is a *trans*-acting regulator of neural stem cell (NSC) differentiation, but the molecular mechanisms by which *Pnky* regulates neurogenesis is unknown. A fundamental step towards mechanistic understanding is to determine whether a lncRNA has folded structure that underlies biological function. Using chemical probing and high-throughput analysis, we determined the secondary structure of *Pnky* folded *in vitro* and *in cellulo*. *Pnky* adopts a compact structure *in vitro* with distinct modules and evidence of tertiary interactions. *In cellulo, Pnky* structure is remarkably similar to the *in vitro* conformation. We used locked nucleic acid oligonucleotides to interrogate the entire *Pnky* transcript for function in NSCs and identified regions that when targeted increased neurogenesis – phenocopying *Pnky* knockdown – without decreasing transcript abundance. Our findings provide a structural basis for the role of *Pnky* in neurogenesis and, more broadly, illustrate how structural maps combined with phenotypic data can advance fundamental understanding of lncRNA mechanism.

**HIGHLIGHTS:**

*Pnky* folds into highly structured conformation *in vitro* with distinct modules.
*Pnky* is a compact lncRNA with evidence of tertiary interactions.
*Pnky* structure in neural stem cells is similar to its *in vitro* conformation.
LNA ASO perturbations identify structured regions of *Pnky* involved in neuronal differentiation.

## Introduction

Neurogenesis – the birth of new neurons from neural stem cells (NSCs) – is a highly orchestrated process that involves the function of numerous genes, some of which “restrain” neuronal production, ensuring that proper numbers of neurons populate the brain across the continuum of development^1,2^. In recent years, a growing number of long noncoding RNAs (lncRNAs) – transcripts longer than 200 nucleotides (nt) that do not code for protein – have been identified as regulators of neurogenesis^3,4^. However, although it is now clear that certain lncRNAs are important to brain development, knowledge fundamental to the understanding of lncRNA molecular mechanism(s) remains limited.

The lncRNA *Pnky* (*Pou3f2* intergenic non-koding) is a nuclear-enriched, evolutionarily conserved neural lncRNA that regulates mouse NSCs both *in vitro* and *in vivo* ^5,6^. In NSCs cultured from mouse brain, either *Pnky* transcript knockdown (KD) or *Pnky* conditional knockout (*Pnky*-cKO) increases neuronal production by up to 5-fold ^5–7^. *In vivo*, genetic deletion of *Pnky* results in abnormal cortical development related to this phenotype of accelerated neurogenesis ^6^, and *Pnky*-null mice exhibit abnormal learning and behavior ^8^. Furthermore, expression of *Pnky* from a bacterial artificial chromosome (BAC) transgenic insertion (BAC-*Pnky*) rescues the *Pnky*-deletion phenotype both *in vitro* and *in vivo* including behavioral abnormality ^6,8^, providing molecular-genetic evidence that this lncRNA functions *in trans*. Thus, multiple orthogonal methods indicate that *Pnky* is a potent, *trans*-acting regulator of neurogenesis that is consequential to neurodevelopment.

A critical knowledge gap in the field of lncRNA biology is that the molecular mechanisms of these transcripts are poorly understood ^9^. In some cases, RNA transcripts fold into secondary and tertiary structures that directly underlie biological function ^10,11^. Thus, determining whether an RNA has defined, folded structure is a fundamental step towards determining molecular mechanism. Although there is a growing number of reports that describe lncRNA structure and provide evidence for their biological function ^11,12^, lncRNAs can also regulate biology through unstructured RNA regions ^13^. Furthermore, for lncRNA genes that function *in cis*, even the act of transcription – independent of RNA sequence – can activate the expression of neighboring genes ^14–16^. Thus, just as distinguishing *cis*-versus *trans*-activity is fundamental to the molecular-genetic characterization of lncRNA gene function ^17^, a logical next step towards mechanistic understanding is to determine whether RNA transcripts from the gene fold into discrete 3-dimensional (3D) structures.

Currently, it is not possible to reliably predict lncRNA structure from sequence alone^18^. To overcome this limitation, chemically probing RNA secondary structure generates experimental data that can drive accurate RNA structure modeling^19–21^. One such method is selective 2′-hydroxyl acylation analyzed by primer extension and mutational profiling (SHAPE-MaP), which has yielded accurate maps of RNA secondary structure in several viral genomes and noncoding RNAs ^22,23^. For insight into RNA tertiary structure, terbium (III) (Tb^+3^)-cleavage analyzed by sequencing (Tb-seq) was recently developed as a high-throughput method for probing transcript for regions that constitute tertiary RNA interactions ^24^. For lncRNAs, such maps of tertiary structure have not been reported.

Knowing the structure of lncRNAs as they exist inside the cell is critically important to understanding molecular mechanism, since the folded conformation of the lncRNA *in vitro* and *in cellulo* could be very different. Most experimentally determined models of lncRNA structure have been developed from studies of transcripts produced *in vitro* ^25–28^. While it is relatively straightforward to study the secondary structure of RNAs *in vitro*, determining the folded state of the RNA inside the cell is far more challenging, particularly for lncRNAs that are of low abundance ^13,29–34^. Thus, although knowledge of the folded state of lncRNAs *in cellulo* is critically important for understanding mechanism, there are only a few examples in the current literature, and none have been reported for lncRNAs that regulate brain development.

In this work, we experimentally determined the structure of *Pnky* transcribed *in vitro* and also as it exists *in cellulo* in the nucleus of NSCs isolated from the mouse brain. We discovered that *Pnky* is a highly compact lncRNA, organized with modules of intricate secondary structure that have evidence of tertiary interactions. The structure of *Pnky in vitro* and *in cellulo* was remarkably similar. We integrated this knowledge of *Pnky* RNA structure with the complex cellular phenotype of neuronal differentiation by overlaying the phenotypic data from LNA oligonucleotide-based experimental perturbations that systematically “tile” across the entire *Pnky* transcript. Together, our results characterize *Pnky* as a lncRNA with potent neurodevelopmental function related to its 3D RNA structure, and more generally, illustrate how a stepwise, systematic approach to lncRNA gene studies can build foundationally to an understanding of the mechanisms of their cellular functions.

## Results

### *Pnky* adopts a homogeneous, compact and stable structure

Many functional non-coding RNAs exert biological effects as compact, well-folded molecules ^11,35–37^. To study the structure of *Pnky* in solution, we used a non-denaturing (or “semi-native”) purification protocol that preserves RNA secondary structure formed during *in vitro* transcription ^38^. With size-exclusion chromatography (SEC), *in vitro* transcribed *Pnky* eluted a single, narrow peak, indicating that this lncRNA exists in solution as a relatively homogeneous population in terms of overall size (**Figure 1A**).

**Figure 1.**
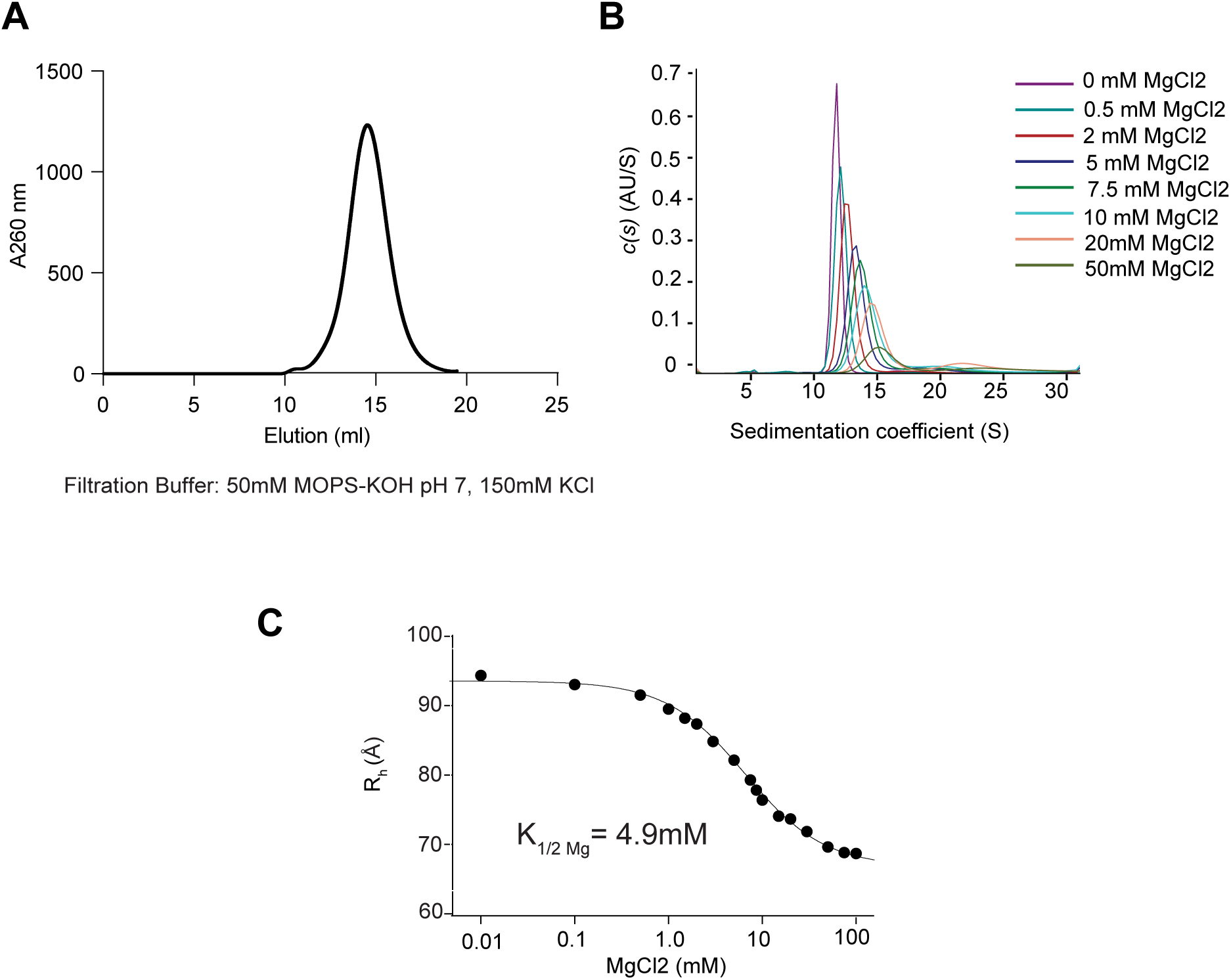
*Pnky* adopts a homogeneous, compact and stable structure. (A) Homogeneity of purified and folded *Pnky* RNA assessed by the size exclusion chromatography (SEC). (B) SV-AUC profiles of *Pnky* RNA obtained with increasing concentrations of magnesium. The graph was obtained using SedFit. (C) Hill plot of the hydrodynamic radii (R_h_, in angstroms) derived from the SV-AUC experiment.

In RNA structure, magnesium ions (Mg^2+^) play an important role in stabilizing RNA folding by neutralizing negative charges on the phosphate backbone ^39^. We performed sedimentation-velocity analytical ultracentrifugation (SV-AUC) at physiological potassium (K+) concentration to monitor the degree of molecular compaction as a function of Mg^2+^ concentration (**Figure 1B**). In SV-AUC analysis, RNA compaction is observed as an increase in sedimentation velocity and a decrease in UV absorption ^40^. As Mg^2+^ concentration increased, the sedimentation coefficient of *Pnky* increased and UV absorption decreased (**Figure 1B**), indicating a magnesium-dependent, global compaction of the *Pnky* molecule. Across the wide range of Mg^2+^ concentrations (0 to 100 mM), *Pnky* remained monodisperse, consistent with this lncRNA adopting a homogeneous, compact structure.

We used SV-AUC data to determine the hydrodynamic radius (R_h_) of *Pnky*. With increasing concentrations of Mg^2+^, *Pnky* RNA compacts with the radius decreasing from ∼ 95Å to 74.4Å. Fitting the R_h_ values at each Mg^2+^ concentration to the Hill equation yielded a K_1/2 Mg_ of 4.9 mM (**Figure 1C**), a value comparable to that of lncRNA *RepA*/*Xist* (K_1/2 Mg_ 4.8 mM) ^41^and even lower than those of other well-structured non-coding RNAs such ai5ɣ group IIB intron (K_1/2 Mg_ 15 mM) ^42^. The low K_1/2_ Mg^2+^ for *Pnky* compaction signifies a high degree of structural stability.

### *Pnky* has intricate secondary structure *in vitro*

To experimentally examine the secondary structure of *Pnky*, we used selective 2′-hydroxyl acylation analyzed by primer extension and mutational profiling (SHAPE-MaP) ^20,22,43^. For SHAPE-MaP, RNA molecules are treated with a hydroxyl-selective electrophile that selectively acylates conformationally flexible nucleotides at the 2’-hydroxyl, resulting in a bulky adduct at single-stranded and/or flexible nucleotides. During MaP readout, the bulky 2’-O-adducts are reported as mutations in the cDNAs produced by reverse transcription (RT).

For *Pnky* SHAPE-MaP, we used 1-methyl-7-nitroisatoic anhydride (1M7) as the electrophile for RNA modification. Based on SV-AUC analysis, we determined that at 15mM Mg^2+^ (three times the experimentally determined K_1/2 Mg_), *Pnky* is fully folded and monodisperse and thus optimal for structural studies. We treated folded *Pnky* with 1M7 or DMSO control, performed MaP-RT and amplified the resultant cDNA with *Pnky*-specific primers, producing two overlapping amplicons that cover the entire length of the RNA (**Figure S1A-B**) for readout by high-throughput sequencing (**Figure 2A**).

**Figure 2.**
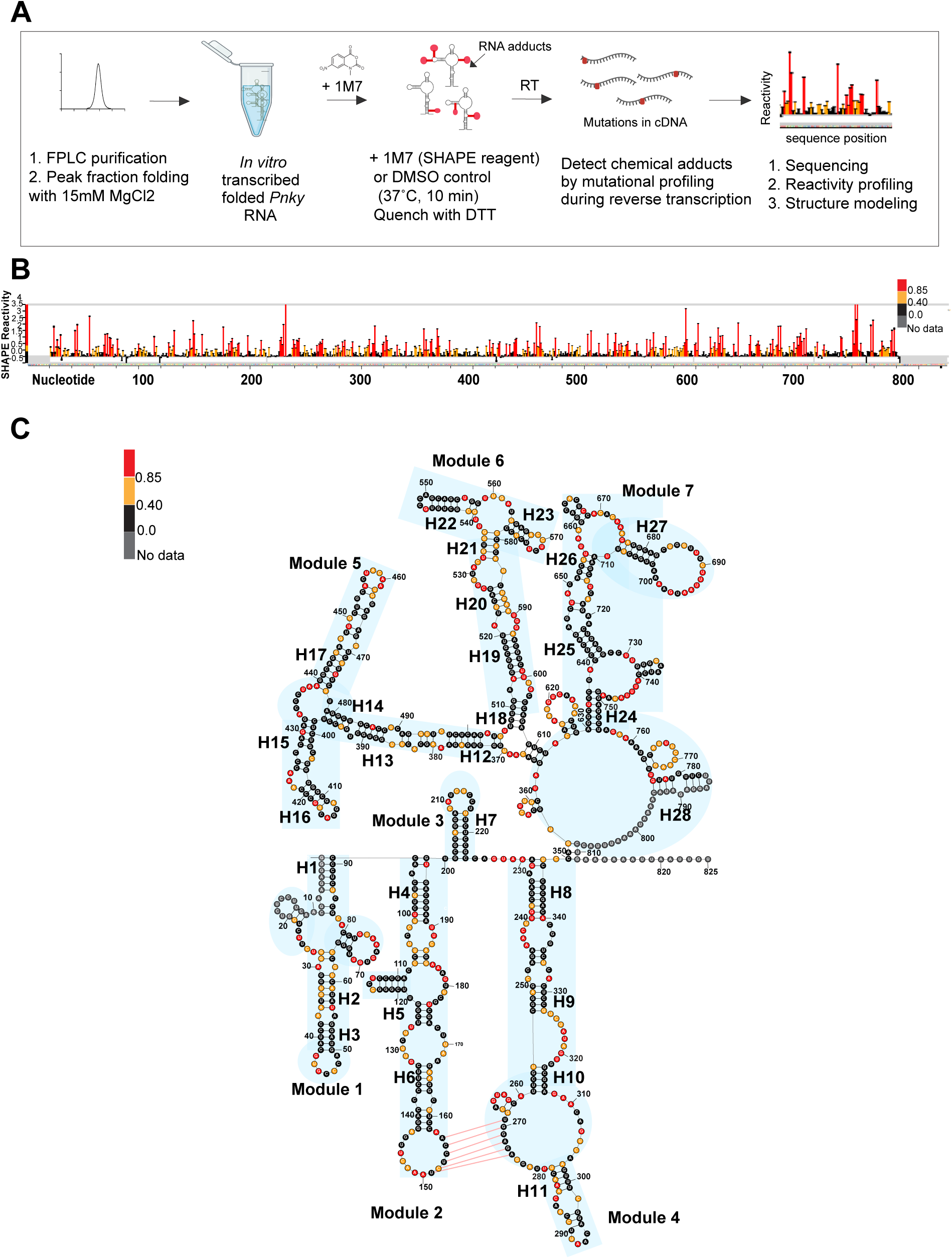
*Pnky* has intricate secondary structure *in vitro*. (A) Schematic of the in vitro SHAPE MaP experiment using in vitro transcribed *Pnky* RNA. (B) SHAPE reactivity profile of *Pnky* RNA at nucleotide resolution. (C) Minimal free energy secondary structure model using SHAPE reactivities as constraints. Structural modules are highlighted in blue. Psuedoknot interaction predicted using SHAPEKnots between stem-loops in module 2 and 4 See also Figure S1 and S2 and Tables S1 and S2.

We performed two independent SHAPE-MaP experiments using independently transcribed, folded preparations of *Pnky* RNA obtained on two separate days. For both 1M7-treated and DMSO control, we obtained two datasets (**Figure S2A)**, each having an average read depth > 50,000x and effective reactivity data for 97.8% (763/780) of the nucleotides (**Table S1**). The relative mutation rates of the 1M7 treated samples were significantly higher than those of the DMSO controls (**Figure S2A**), with excellent concordance across the replicates (Pearson correlation r = 0.94) (**Figure S2B**). We used these data to determine SHAPE reactivity at single nucleotide resolution across the length of *Pnky* (**Figure 2A, Figure S2C**).

The SHAPE reactivity data were then used as constraints for *de novo* secondary structure prediction. *Pnky* exhibited highly complex, intricate secondary structure *in vitro*. More than 65% of the nucleotides were involved in base pairing, forming 28 helical segments, 18 terminal loops, 15 internal loops, and 12 junction regions (**Figure 2C, Figure S2D**), which is comparable to other noncoding RNAs known to be highly structured ^44^.

Examination of the secondary structure map revealed seven distinct structural modules (highlighted in blue), each formed of multiple helical segments, loops, and junction regions. (**Figure 2C**). Module 1 (nt 1-91) has three helices (H1-H3), one 4-way junction and three hairpin stem loops. Module 2 (nt 92-200) consists of three helices (H4-H6) with internal loops, a three-way junction, and terminal hairpin stem loop. Module 3 (nt 201-224) consists of a single helical region (H7) with a hairpin loop. Module 4 (nt 232-350) has 4 helices (H8-H11), a short single-stranded region (nt 225-231), and a three-way junction. Modules 5 and 6 (nt 371-505 and nt 507-608, respectively) each consist of six helical regions (H12-H17 and H18-H23, respectively) that fold into dumbbell-like structures. Both Modules 5 and 6 radiate from a short stem that emerges from a wheel-like structure of Module 7 (nts 613-774) from which five other substructures containing five helices (H23-H28) emanate.

RNA helices can assemble into a tertiary structure called a “pseudoknot,” which consists of two stem-loop structures in which half of one stem is interposed between the two halves of another stem. RNA pseudoknots play important roles in the expression of many genes ^29,37,45^. Using ShapeKnots ^46^, a pseudoknot was predicted at terminal loops of Modules 2 and 4 (indicated by red lines in **Figure 2C**), with base-pairing interactions across the single stranded regions of the loops being supported by low SHAPE reactivities at nucleotides 151-156 and 269-274 (**Figure 2B, Figure S2C**). Furthermore, incorporating the pseudoknot as a hard constraint in structure prediction lowers the Shannon entropy, suggesting that the pseudoknot-containing model is more thermodynamically stable (**Figure S2E**).

### Tb-seq detects regions of tertiary structure in *Pnky*

The highly compact, stable and monodisperse nature of *Pnky* in solution is a strong indication that this lncRNA contains specific tertiary structures that underlie its overall 3D structure. Identifying regions of stable RNA tertiary structure is challenging, which likely explains why very few lncRNAs have been studied at this level.

To form specific tertiary interactions, multiple RNA secondary structures must bring their negatively charged phosphate backbones into close proximity. Such interactions are often stabilized by the local coordination of multivalent cations such as Mg^2+^, which neutralize local negative charge. Lanthanide ions, such as terbium (III) (Tb^3+^), can take the place of Mg^2+^ within an RNA structure, but due to its chemical nature, Tb^3+^ induces local RNA cleavage at the site of binding. Recently, a high-throughput sequencing method, Tb-seq, was developed to identify sites of Tb^3+^ cleavage, systematically identifying sites of tertiary interactions within an RNA molecule^24^.

We used Tb-seq to study the tertiary structure of *Pnky*. Briefly, after treating *in vitro* transcribed, folded *Pnky* with Tb^3+^, we performed RT, and sites of strand cleavage were identified with high-throughput sequencing (**Figure 3A**). For each nucleotide, we determined the probability of Tb^3+^-dependent cleavage vs. spontaneous cleavage, generating a reactivity score (see Methods) (**Figure S3A)**. The reactivity scores across two independent replicates were highly similar (Pearson correlation r = 0.97) (**Figure S3B**).

**Figure 3.**
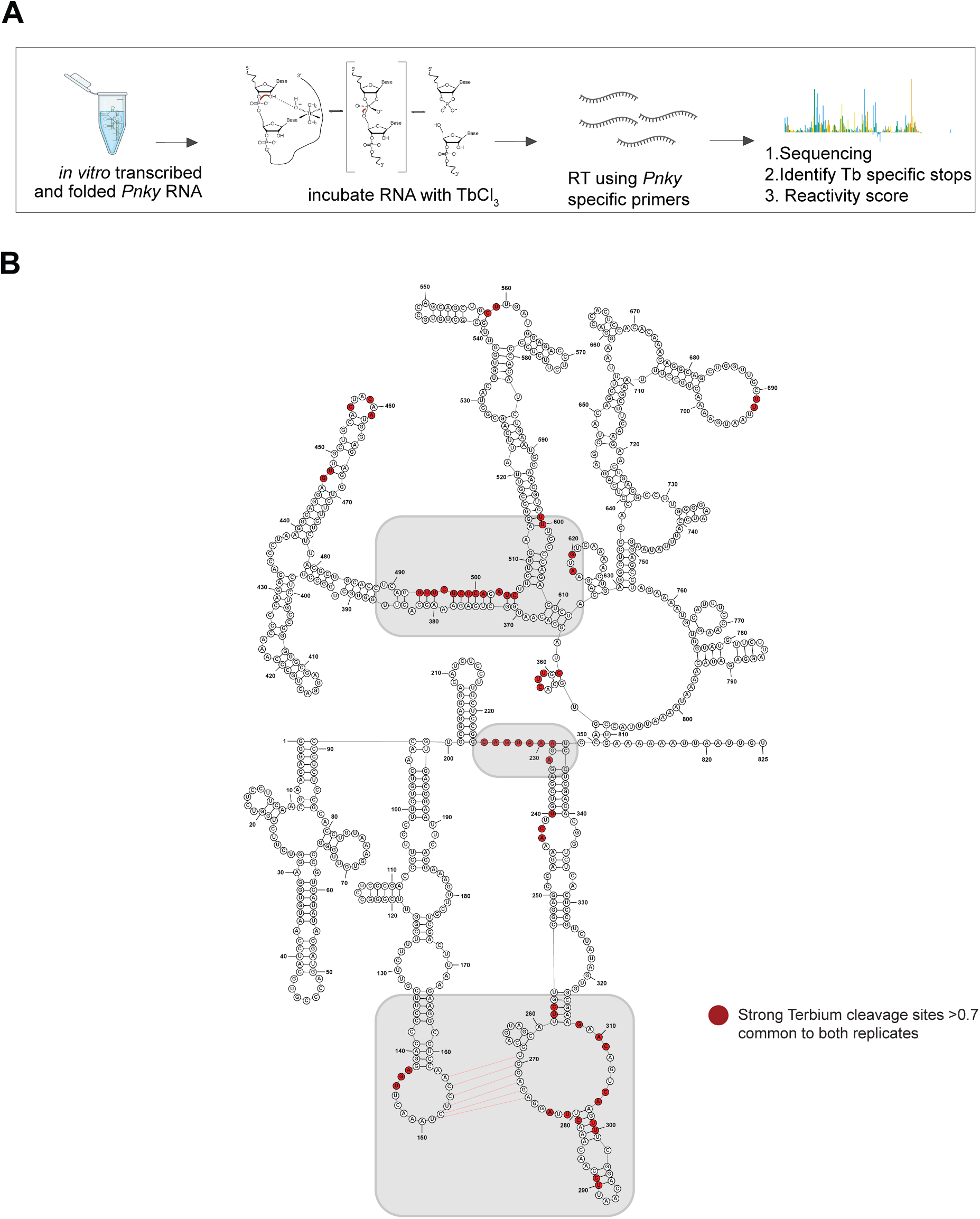
Tb-seq detects regions of tertiary structure in *Pnky*. (A) Schematic showing Terbium-sequencing workflow. (B) Secondary structure of the *Pnky* RNA displaying sites of strong Tb^3+^cleavage (red) with three regions of concentrated Tb reactivity highlighted in grey. See also Figure S3 and Table S1.

To identify sites that likely result from specific, site-bound Tb^3+^-dependent cleavage, we used two criteria ^24^. First, a stringent reactivity value of >0.7 was used as a threshold for each site. Second, these sites must be observed in two independent replicates. Nucleotide sites that satisfy these criteria are highlighted in red in the secondary structure diagram of *Pnky* (**Figure 3B, Figure S3C**).

*Pnky* had 57 sites of Tb^3+^ induced cleavage. The distribution of most (70.2%) Tb^3+^ cleavage sites was concentrated in three major regions of the *Pnky* secondary structure model (**Figure 3B**). One region corresponds to the predicted pseudoknot that crosses Modules 2 and 4, with Tb^3+^ reactivity prominent around the “kissing loop” component (**Figure 3B**, highlighted in grey). The second major region is located at the boundary between Module 3 and 4 (**Figure 3B**, grey box). The third region of Tb^3+^ reactivity is found among the nucleotides of Modules 5, 6 and 7 that map to secondary structures that are proximal to their emanation from the major ring of Module 7 (**Figure 3B**, grey box). In addition to these three clusters of Tb^3+^ reactivity, a minor number of sites (29.8%) are found among nucleotides of distal aspects of secondary structures of Modules 5, 6 and 7. Together, these data represent the first systematic analysis of tertiary structure in a lncRNA, and for *Pnky*, provide experimental evidence for a specific 3D conformation.

### *Pnky* structure *in cellulo* is similar to its *in vitro* folded state

It is still fundamentally unclear whether *Pnky* has a defined structure *in cellulo* that underlies its biological function. Thus, despite having a high quality *in vitro* structural map, it is also important to obtain a parallel map that captures the conformational states of *Pnky in cellulo*. Furthermore, knowledge of *Pnky* structure *in vivo* is important to the interpretation of experiments that interrogate the potential biological function of specific regions of the lncRNA.

SHAPE-MaP can be performed *in cellulo* by treating cells with the electrophile reagent, extracting the RNA, then performing RT and downstream sequencing analysis ^47^. For RNAs that are abundant, such as the genome of RNA viruses, *in cellulo* SHAPE-MaP has provided data of sufficient depth for *de novo* structural modeling ^23^. However, for RNAs that are at low abundance, such as most lncRNAs, *in cellulo* SHAPE-MaP reactivity data has been mostly used to corroborate selected regions of the *in vitro* conformation, not to generate an entirely structural model *de novo* ^29^.

We sought to perform SHAPE-Map for *de novo* secondary structural modeling of *Pnky* as it exists in the cellular nucleus of primary neural stem cells that exhibit a strong *Pnky*-dependent phenotype rather than in the immortalized cell lines. Given our prior results ^5,6^ we selected NSC cultures established from the postnatal ventricular subventricular zone (V-SVZ) for our *in vivo* SHAPE-MaP studies.

*Pnky* is nuclear-enriched lncRNA that is at relatively low abundance, approximately 10-20 copies per nucleus, which presents an important technical challenge. For *in cellulo* SHAPE-MaP to be successful, it is necessary to acylate the target RNA inside the cell, which is far less efficient than for purified RNA *in vitro*, and the low concentration of *Pnky* necessitates the use of relatively large numbers of cells cultured from a specific part of the mouse brain. In early experiments, we used 70-90 million V-SVZ NSCs cultured from postnatal day 7 (P7) *Pnky^+/+^* mice for *in cellulo* SHAPE-MaP, however SHAPE reactivity data was not sufficient for reliable *de novo* structural modeling.

Increasing RNA transcript abundance can improve *in cellulo* SHAPE-MaP studies ^48^. Although it is possible to express *Pnky* at very high levels from lentiviral vectors ^49^, we have found that vector-expressed *Pnky* transcripts localize primarily in the cytoplasm (unpublished observations), unlike the normal nuclear localization of *Pnky* ^5^. In contrast, BAC-expressed *Pnky* – which rescues *Pnky*-KO phenotypes including at the level of the transcriptome – remains nuclear-enriched ^6^; the levels of *Pnky* in V-SVZ NSC cultures from *Pnky*^+/+^; BAC-*Pnky* mice were approximately 10-12-fold higher than those from *Pnky*^+/+^ mice.

Thus, for *Pnky in cellulo* SHAPE-MaP analysis, we used NSC cultures established from the V-SVZ of P7 *Pnky*^+/+^; BAC-*Pnky* mice. For each biological replicate, we established primary V-SVZ NSC cultures from multiple P7 *Pnky*^+/+^; BAC-*Pnky* litters, treated 70-90 million NSCs with either 1M7 (50mM) or DMSO, isolated the nuclei, extracted nuclear RNA, then depleted ribosomal RNA to enrich for *Pnky* RNA (**Figure 4A**). After RT, PCR amplification with *Pnky*-specific primers produced two overlapping amplicons of expected size (**Figure S4A**), generating sequencing libraries that cover the entirety of the 825-nucleotide transcript. To the best of our knowledge, this is the first *in-cell* SHAPE data for a neural RNA from the primary cell culture of neural stem cells. Across the two replicates, we obtained effective reactivity data for 97.8% of the nucleotides (**Table S1**) and significantly higher mutation rates with 1M7 as compared to DMSO control, indicating successful modification of the RNA *in cellulo* (**Figure S4B**). Furthermore, SHAPE reactivities from both replicates had strong correlation across the entire *Pnky* transcript (Pearson correlation r = 0.84) (**Figure 4B, Fig S4C-D**).

**Figure 4.**
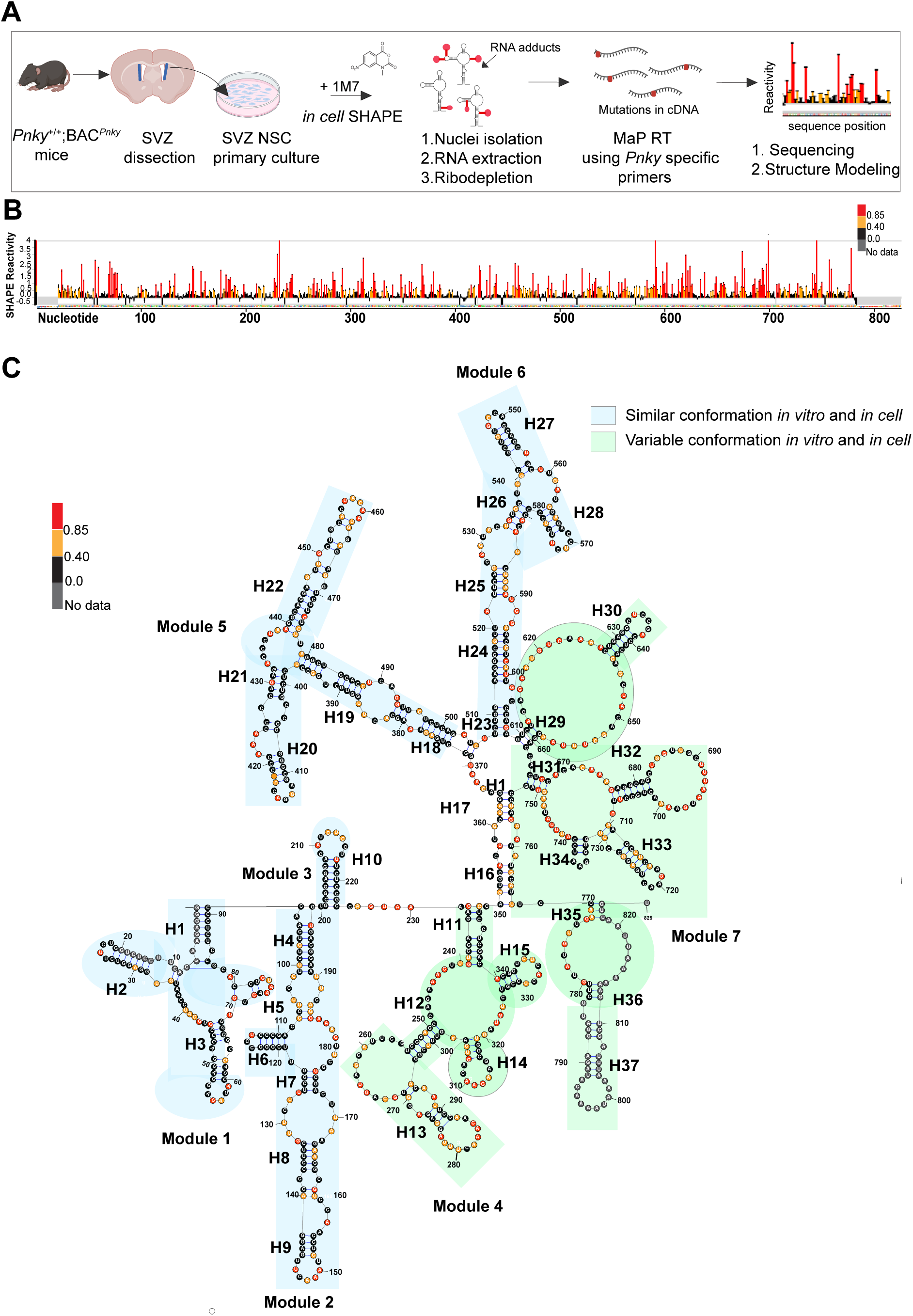
*Pnky* structure *in cellulo* is similar to its *in vitro* folded state. (A) Schematic of the *in-cell* SHAPE MaP for *Pnky* RNA in primary culture of mouse neural stem cells. (B) *In-cell* SHAPE reactivity profile of *Pnky* RNA at nucleotide resolution (C) Minimal free-energy secondary structure model using *in-cell* SHAPE reacitivites at nucleotide resolution for *Pnky* RNA. Blue and green shaded boxes show the modules with similar and variable conformations *in vitro* and *in cell* structural models See also Figure S4 and Tables S1 and S2.

We then used *in cellulo* SHAPE reactivities to experimentally constrain secondary structure modeling ^50^. Nearly 70% of the nucleotides were involved in base pairing, forming 37 helical segments, 18 terminal loops, 18 internal loops, and 10 junction regions (**Figure 4C, Figure S4E**), which is similar to the degree of structure observed for *in vitro Pnky* (**Fig 2C**). Thus, like its folded state *in vitro, Pnky in cellulo* is highly structured with a similar distribution of RNA structural motifs.

Like the conformation of *in vitro Pnky* (**Figure 2C**) the secondary structure of *in cellulo Pnky* can be divided into seven modules (**Figure 4C**). Across the two structural models – *in vitro* and *in cellulo*, both developed *de novo* and thus independent from one another – the number of nucleotides within each module is very similar, and the borders between each module are within few nucleotides of one other (**Table S2**).

While five *Pnky* modules have similar secondary structural organization across the two models, two modules (Modules 4 and 7) appear different *in cellulo* as compared to *in vitro*. *In vitro*, Module 4 emerges after a short single stranded region that builds into a structure containing three internal bulges, the last of which is predicted to interact with a terminal loop of Module 2. *In cellulo*, Module 4 contains three internal bulges that have a different arrangement, and the pseudoknot interaction across Modules 4 and 2 are not predicted. Module 7 exhibits even greater differences across the i*n cellulo* and *in vitro* models. Whereas *in vitro* Module 7 emerges mostly from a wheel-like structure, *in cellulo*, this central wheel is not present. These data illustrate potential differences in secondary and tertiary structure *in cellulo* as compared to its folded state *in vitro*.

Five of the seven modules of *Pnky* are remarkably similar across the *in cellulo* and *in vitro* models. Such similarity can result from these regions of RNA folding equivalently in the two different environments and if these regions *in cellulo* do not strongly interact with other macromolecules such as proteins ^51^. Across the two models, each corresponding module shares a similar overall shape and pattern of secondary structural motifs. For example, both *in cellulo* and *in vitro*, Module 1 can be characterized by an internal bulge from which three hairpin loops emerge. In both models, Module 2 consists of a helix followed by an internal bulge, followed by a bulge with a side-protruding hairpin loop, then another bulge from which another structure emerges that ends in a terminal hairpin loop. Module 3 is nearly identical, being defined by a solitary hairpin loop. Modules 5 and 6 are dumbbell shaped and have highly similar patterns of helices, internal bulges and hairpin loops. Thus, these models of *Pnky* structure *in cellulo* and *in vitro* – both of which were developed *de novo* and independent of one another – exhibit remarkable similarity with distinct, focal regions of difference, providing information critical for understanding the biological function of this lncRNA inside the cell.

### Specific LNA ASOs that target *Pnky* can phenocopy *Pnky*-KD

Disrupting RNA secondary structures with LNA antisense oligonucleotides (ASOs) is a powerful approach for determining if observed RNA structures are in fact functional ^52,53^. LNAs are non-natural RNA base analogs that increase the melting temperature (T_m_) of nucleoside duplexes, which allows LNA ASOs to outcompete endogenous RNA-RNA duplexes that comprise secondary structure ^54^. Because LNAs have non-natural chemical structure, the binding of LNA ASOs does not activate RNase H-mediated degradation of the RNA target ^54^. Thus, LNA ASOs have the potential to identify lncRNA function related to RNA structure and not simply lncRNA transcript abundance.

To evaluate the *Pnky*-targeting LNA ASOs for function in neural development, we used the well-characterized primary V-SVZ NSC culture system that recapitulates key aspects of neurogenesis *in vivo* ^55–57^. V-SVZ NSC cultures can be passaged as a readily transfectable monolayer, and neuronal differentiation is induced by removal of mitogenic factors (**Figure 5A**). In V-SVZ NSC cultures, *Pnky* transcript KD or *Pnky* cKO has a “positive” phenotype, increasing neurogenesis ^5,6^.

**Figure 5.**
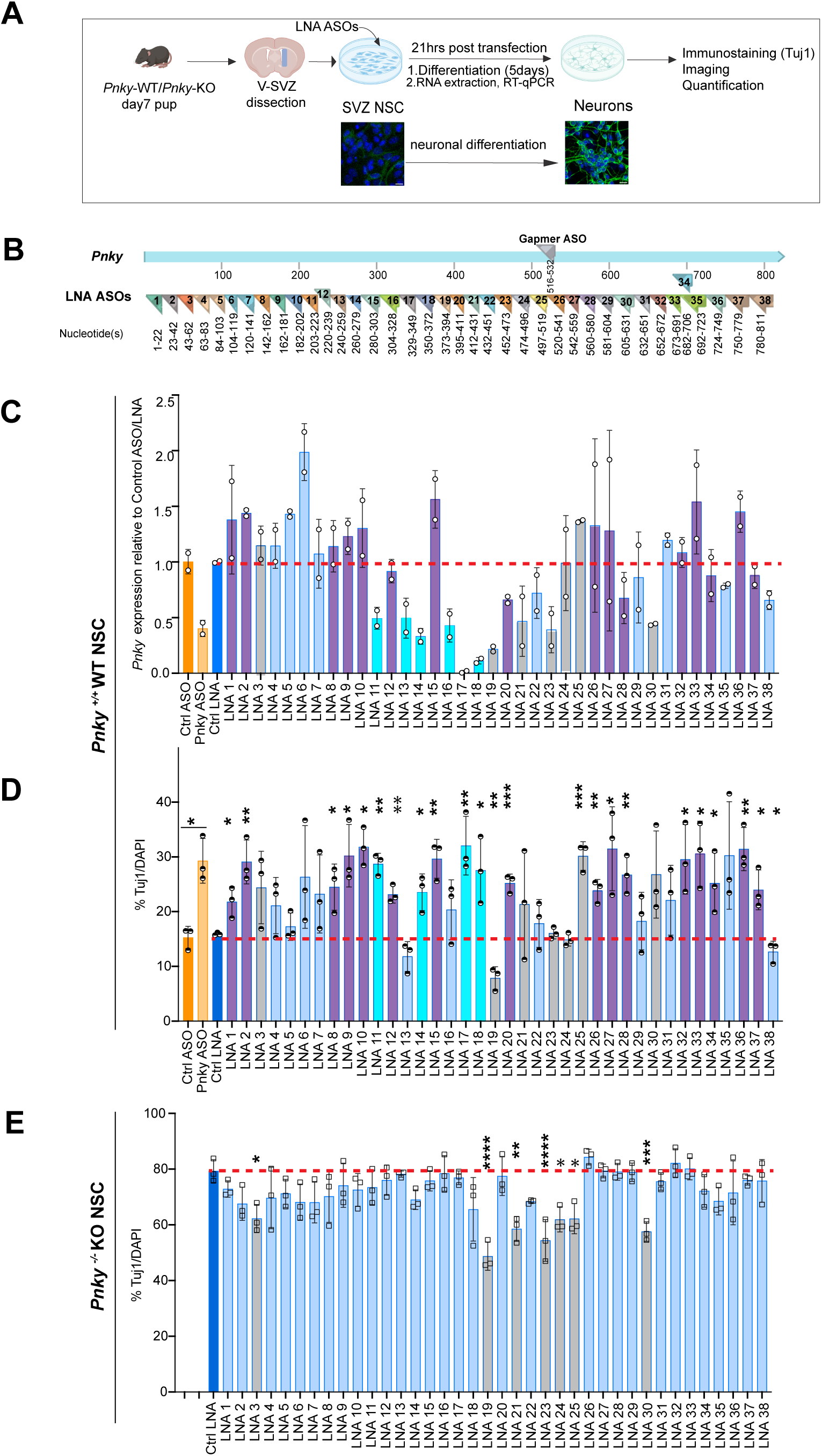
Specific LNA ASOs that target *Pnky* can phenocopy *Pnky*-KD. (A) Schematic of the LNA and Gapmer ASO transfection in the NSCs and differentiation assay (B) Location of LNA and gapmer ASOs binding sites on the *Pnky* RNA. (C) Relative fold-change in *Pnky* transcript levels in *Pnky-*WT V-SVZ cultures by RT-qPCR 21-24 hours after transfection with LNA and Gapmer ASOs. (D) Tuj1 ICC in d5 differentiated V-SVZ cultures established from *Pnky*-WT mice. Quantification using % Tuj1 normalized to DAPI positive cells in *Pnky*-WT NSCs. (E) Tuj1 ICC in d5 differentiated V-SVZ cultures established from *Pnky*-KO mice. Quantification using % Tuj1 normalized to DAPI positive cells in *Pnky*-KO NSCs. \**p* < 0.05 \*\**p* < 0.01, \*\*\**p* < 0.001, \*\*\*\**p* < 0.0001, ns = non-significant. Data represented as mean+/ SD for n= 3 independent technical replicates of V-SVZ culture obtained from pooled dissection of postnatal day d7 animals. Two-tailed unpaired t-test for WT dataset and Ordinary one-way ANOVA with multiple comparisons for the KO dataset. See also Figures S5 and S6 and Tables S3.

We used LNA ASOs to interrogate the length of the *Pnky* transcript for biological function. Instead of targeting selected *Pnky* secondary structures predicted by SHAPE-MaP analyses, we chose to pursue an unbiased approach by designing a set of 38 LNA ASOs that “tile” the entire *Pnky* lncRNA (**Figure 5B, Table S3**). Establishing a rigorous and scalable neuronal differentiation assay for LNA ASO studies is challenging given the developmental complexity of neurogenesis, which can be adversely affected by experimental interventions such as transfection. To benchmark the assay and illustrate the positive neurogenic phenotype of *Pnky*-KD in LNA transfection studies, we produced an LNA ASO with a “gapmer” design, wherein the central nucleotides of the oligonucleotide are chemically unmodified DNA, which allows RNase H-mediated degradation of the RNA target ^52,54^. One day (1d) after transfection, the *Pnky* gapmer ASO reduced *Pnky* transcript levels by ∼55% as compared to non-targeting gapmer control (**Figure 5C**, orange bars), and after 5 d of differentiation, gapmer-mediated *Pnky*-KD increased neurogenesis by nearly 2-fold (**Figure 5D**, orange bars), as expected.

We assayed in parallel all 38 LNA ASOs alongside gapmer ASO and control oligonucleotides in triplicate NSC cultures from *Pnky* wild-type (WT) mice to assess the neuronal production phenotype (**Figure 5D, Figure 5SA-B**). As compared to transfection of LNA oligonucleotide control (**Figure 5D**, dark blue bar), 21 LNA ASOs that target *Pnky* increased neuronal production by 1.5-to 2-fold (**Figure 5D**, purple bars and a subset of grey bars) and one LNA ASO reduced neurogenesis by 0.9-fold (**Figure 5D**, LNA 38). Thus, overall, 58% (22 of 38) of the LNA ASOs produced a neurogenic phenotype, and of these, the vast majority (95%, 21 of 22) phenocopied the effect of *Pnky*-KD.

To filter for LNA ASOs with potential off-target effects, we also assayed in parallel all 38 LNA ASOs in NSC cultures from *Pnky*-KO mice (**Figure 5E, Figure S6A-B**). None of the LNA ASOs increased neurogenesis from NSCs that lack *Pnky* transcript. However, we did identify 7 LNA ASOs that reduced neuronal production in *Pnky*-KO NSCs (**Figure 5E**, grey bars). Thus, while 7 LNA ASOs had a *Pnky*-independent negative effect upon neurogenesis, none of the other 31 LNA ASOs had evidence of an off-target positive neurogenic phenotype.

Although LNA ASOs do not directly induce degradation of the RNA target, loss of the RNA target could still be observed after transfection. We therefore next filtered for LNA ASOs that reduced *Pnky* levels after transfection, since such a change would confound the interpretation of a neurogenic phenotype and its potential relationship to RNA structure. Among the 31 LNA ASOs without evidence of off-target effect, 6 reduced the detection of *Pnky* to less than 50% of control (**Figure 5B**, LNAs 11, 13, 14, 16, 17, and 18). Of note, 4 of these 6 LNA ASOs (LNAs 11, 14, 17, 18) increased neurogenesis by 1.5 to 2.1-fold (**Figure 5C**, teal bars), similar to the phenotype observed with gapmer-mediated *Pnky*-KD (**Figure 5C**, orange bars).

To address the fundamental question of whether *Pnky* can regulate neural development through its RNA structure independent of overall transcript abundance, we examined the 25 LNA ASOs that did not reduce *Pnky* transcript levels or have off-target effects. In this filtered subset, 28% (7 of 25) of the LNA ASOs did not increase neurogenesis (**Figure 5D**, light blue bars), indicating that the transfection of *Pnky*-targeted LNA ASOs does not generally promote neurogenesis. Importantly, however, 72% (18 of 25) of the LNA ASOs increased neuronal production by 1.5 to 2.1-fold (**Figure 5D**, purple bars). The identification of multiple *Pnky*-targeted LNA ASOs that increase neurogenesis without decreasing *Pnky* transcript levels provides foundational evidence for RNA structure-based mechanism of this neurodevelopmental lncRNA.

### LNA ASO phenotypic data maps to specific structured regions of *Pnky*

To investigate whether the structural maps of *Pnky* provide insight into its biological function, we mapped the location of all 38 LNA ASOs onto the models from *in cell* (**Figure 6A**) and *in vitro* SHAPE-MaP data (**Figure 6B**). The seven LNA ASOs (locations shown in grey) that had evidence of off-target effects were excluded from further analyses. The locations of the six LNA ASOs that led to reduced levels of *Pnky* were mapped in teal, and because the effects of these LNAs upon neurogenesis are entangled with the known phenotype of *Pnky*-KD, we did not focus attention upon those regions. The locations of the seven LNAs that did not reduce *Pnky* transcript level nor increase neurogenesis are shown in light blue; these negative results indicate that the transfection of LNAs in general does not increase neurogenesis from our NSC cultures.

**Figure 6.**
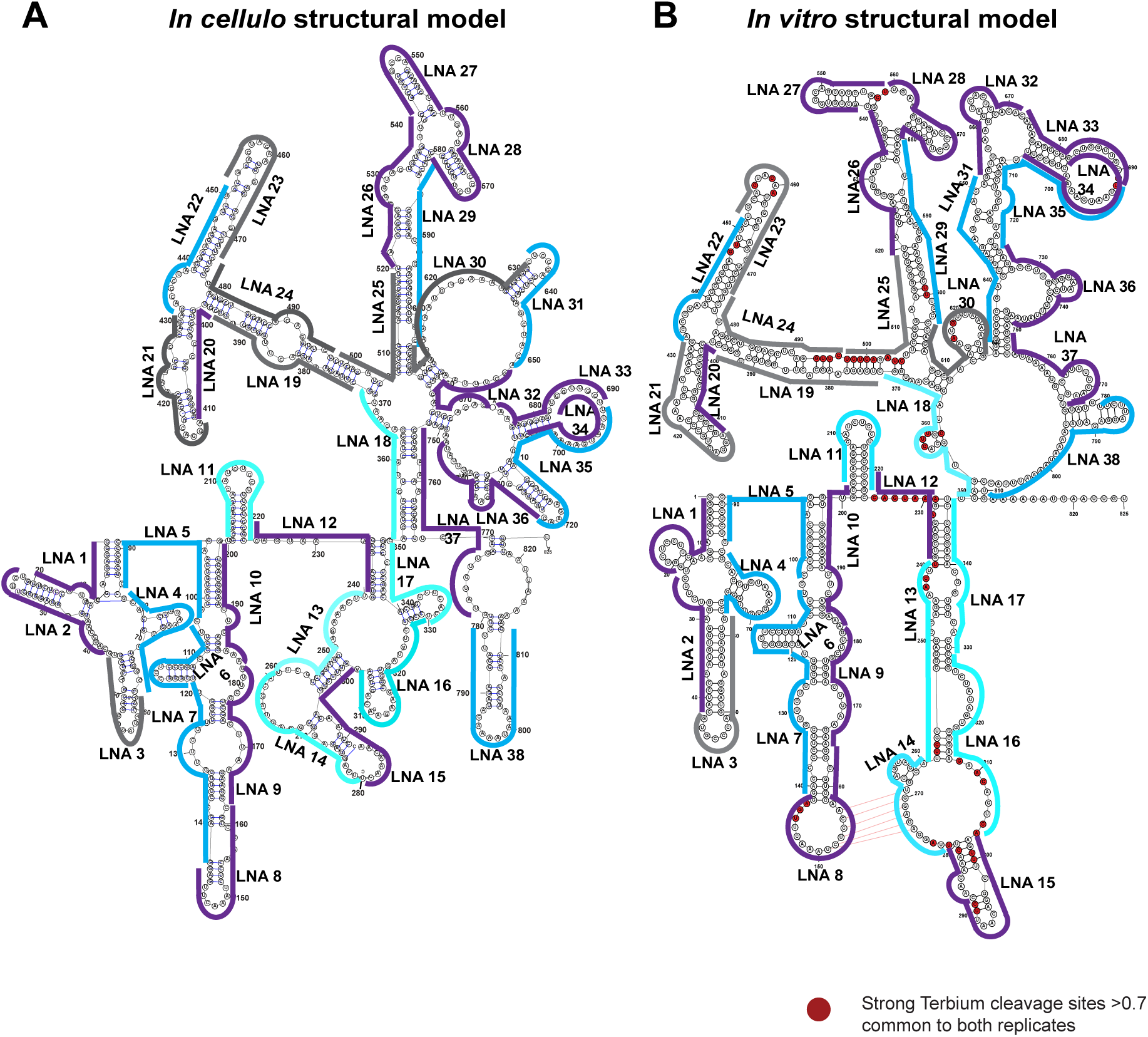
LNA ASO phenotypic data maps to specific structured regions of *Pnky*. (A) maps the locations of the LNA ASOs on the *in-cell* structural maps of Pnky RNA. (B) maps the locations of the LNA ASOs on the *in vitro* structural maps of Pnky RNA. ‘*Pnky*-independent toxic’ LNAs in gray, ‘no phenotype’ LNAs in light blue, ‘destabilizing’ LNAs in teal and ‘increased neurogenesis without decreasing *Pnky* transcript levels’ LNAs in purple. Terbium reactive nucleotides are shown in the *in vitro* structural model in red dots. See also Table S3.

We mapped the locations of the 18 LNA ASOs that increased neurogenesis without decreasing *Pnky* transcript levels in purple. LNA 1 and LNA 2 both target Module 1 toward the 5’ end, and in the *in vivo* model, they map to opposite sides of the first stem-loop. LNAs 8, 9 and 10 map to regions of Module 2 that are remarkably similar across the two models, an RNA segment containing two internal loops, and a distal stem-loop. LNA 8 targets one-half of the pseudoknot predicted *in vitro*, and this region also contains three Tb-reactive nucleotides, indicating that LNA 8 targets a region with evidence of tertiary structure.

LNA 12 targets a region that lies between Modules 3 and 4 in both the *in cellulo* and *in vitro* models. This region contains a stretch of eight Tb-reactive nucleotides and lies between the solitary stem loop of Module 3 and the first internal bulge of Module 4, suggesting a focus of tertiary structure.

LNA 15 targets Module 4, which is conformationally different in terms of secondary structure across the two models. However, it is interesting to note that LNA15 targets a region of local Tb-reactivity, with nine Tb-reactive nucleotides either directly under this LNA or immediately adjacent. Thus, like LNAs 8 and 12, LNA 15 targets a region with evidence of tertiary structure.

Modules 5 and 6 had overall secondary structure similar across the two *Pnky* models, both forming dumbbell-shaped structures in these regions. LNA 20 targets one arm of the dumbbell of Module 5. In Module 6, LNA 26 targets a region immediately proximal to the dumbbell, and LNAs 27 and 28 target the two arms.

Because the overall structure of Module 7 is different *in cellulo* as compared to its *in vitro* conformation, the secondary structural motifs targeted by LNAs 32, 33, 34, 36 and 37 are variable across the two models. Nevertheless, we note that LNAs 33 and 34 are overlapping (due to a redundancy in our LNA targeting design), indicating that targeting the same region of *Pnky* with slightly different LNA reagents can produce the same positive neurogenic phenotype.

In summary, the phenotypic data of these 18 LNAs (mapped in purple in **Fig 6A-B**) identified 17 distinct regions of *Pnky* that are involved in the regulation of neurogenesis from mouse NSCs. Six of the seven modules had at least one LNA that increased neuronal production, suggesting that the function of *Pnky* in neurogenesis is distributed broadly across the lncRNA. The LNAs that target Modules 1, 2, 5 and 6 – structures that are similar *in vitro* and *in cellulo* – indicate that regions that can fold analogously independent of nuclear factors are important for *Pnky* function. LNAs 8, 12 and 15 target regions with evidence of tertiary structure, which prioritizes these modules and potential interactions for involvement in higher-order 3D conformation. The overall secondary structure of Modules 4 and 7 is distinct *in cellulo* as compared to *in vitro*, which can be explained by these regions interacting with other factors in the nucleus ^51^. Thus, LNAs that map to Modules 4 and 7 are particularly interesting, as they may target regions of *Pnky* that interact with regulatory proteins in the cell. Overall, the integration of RNA structural maps with a systematic approach to LNA targeting provides the groundwork from which a structure-function based mechanistic understanding of *Pnky* can be further built.

## Discussion

As compared to other tissues and organs, the mammalian brain is particularly enriched in lncRNAs ^58^, with many of these noncoding transcripts being cell-type specific and dynamically expressed during neurodevelopment ^59–61^. In terms of biological function and molecular mechanism, lncRNAs remain a relatively mysterious class of noncoding transcript. For the growing number of individual lncRNAs that have been shown to regulate neurodevelopment, the molecular underpinning of their function is poorly understood ^4,9^. For our investigation of *Pnky*, we have taken a stepwise, systematic approach to understanding its function in neurogenesis, which has led us to the RNA structure studies presented here.

After using multiple molecular-genetic approaches to establish that *Pnky* is a *trans*-acting lncRNA gene ^6,8^, determining whether the *Pnky* transcript itself adopts a defined 3D structure, particularly *in cellulo*, was a key next step towards understanding its molecular mechanism. Not all lncRNA genes produce transcripts that require specific nucleotide sequences or structured regions for their biological function ^9,11,17^. Thus, although the genetic evidence for *Pnky* function in *trans* suggested that specific RNA structure plays a role in the function of this lncRNA, it was equally plausible that *Pnky* exists as an unstructured transcript both *in vitro* and *in cellulo*. An answer to this question would be critical to generating additional hypotheses regarding its molecular mechanism.

*In vitro*, *Pnky* exhibited a homogeneous, compact structure with remarkable stability. The stability of *Pnky* folding, as assessed by K_1/2 Mg_, is comparable to the RepA region of *XIST* and even lower than that of ai5ɣ group IIB intron, both being examples of RNA structure that appears to play a role in their cellular function ^41,42^. The biophysical stability of *Pnky in vitro* adds importance to the hypothesis that this lncRNA transcript possesses biological information in its 3D structure.

Underlying the stable global conformation of *Pnky in vitro* was an intricate secondary structure comprised of seven modules. Other studies of *in vitro* transcribed lncRNAs have indicated a modular nature of lncRNA secondary structure. For example, *HOTAIR* and the *RepA* region of XIST fold into modular domains that facilitate interaction with chromatin regulators ^25,34,62^. A G-rich internal loop motif in lncRNA *Braveheart* physically interacts with and antagonizes a specific transcription factor critical for cardiomyocyte differentiation ^27^. Thus, like other known functional lncRNAs, *Pnky* appears to be among those that possess intricate, modular structure *in vitro*.

For *Pnky*, a lncRNA that localizes primarily to the nucleus, it was unclear if its SHAPE reactivities *in cellulo* would be vastly different from those observed *in vitro*. Across five of the seven modules in *Pnky*, SHAPE reactivities were remarkably similar *in cellulo* as compared to *in vitro*. Furthermore, these five modules exhibited very similar overall secondary structural patterns in both *de novo* models of the entire transcript. This similarity suggests that the structures of these modules are likely independent of their interaction with nuclear proteins or nucleic acids. The similarity further suggests that the RNA splicing events that occur *in cellulo* do not alter the folding of these five modules.

Modules 4 and 7 exhibited different conformations *in cellulo* as compared to *in vitro*. The fact that the five other modules were structurally similar across the models highlights Modules 4 and 7 as regions of potential biological interactions in the nucleus. The nature of such interactions is not addressed by our studies, and we recognized that it is also possible that RNA splicing *in cellulo* selectively affects Modules 4 and 7 but not the five others. Notwithstanding these limitations, the comparison of *in cellulo* and *in vitro* RNA structural models – both generated de novo from SHAPE reactivity constraints determined with the same chemical probe – determine *Pnky* to be a lncRNA with dense secondary structure (>70% nucleotides base-paired) wherein two of the five modules are highlighted as regions of potential biological interactions in the nucleus of NSCs.

There is very little knowledge about lncRNA tertiary structure. For a very limited number of lncRNAs, techniques such as atomic force microscopy (for *Meg3*)^29^ and small X-ray angle scattering and NMR (for *Bvht*^28^*, XIST*^63–65^) have been useful to gain an overview of the higher order folding. Tb-seq was developed as a comparatively high-throughput method of identifying nucleotides involved in the compaction of RNA into tertiary structures ^24^.

Our Tb-seq analysis of *Pnky* characterized this lncRNA as one with discrete tertiary structure. The focal regions of terbium cleavage highlight regions of *Pnky* that are more likely to play key roles in overall compaction and 3D conformation of secondary structures. Pseudoknots are a tertiary structure that can underlie the function of RNA molecules, and the lncRNA *MEG3* contains conserved pseudoknots that regulate the stimulation of p53 pathway^29^. Although the terbium cleavage sites local to the predicted pseudoknot (**Figure 3B**) provided experimental evidence for this tertiary interaction *in vitro*, we note that the pseudoknot was not predicted in the *de novo* model of *Pnky in cellulo.* The putative pseudoknot in *Pnky* involves an interaction between Modules 2 and 4, and while Module 2 is similar across the two models, Module 4 is different. One intriguing possibility is that *in cellulo,* the *Pnky* pseudoknot is a transient structure, serving to recruit other factors, which subsequently alter the structure of Module 4 through direct binding and/or changes to its conformation.

Studies of lncRNA structure can benefit from comparisons to orthologs, and *Pnky* is an evolutionarily conserved lncRNA gene. For this study, we did not use the strategy of evolutionary comparison because in mice, *Pnky* has been characterized as a fully spliced transcript, whereas the human *PNKY* transcript contains introns between the three conserved exons ^5^. The biological implication of this difference in the linear structure of this lncRNA across these two species is not yet clear, and there are currently no reports of the function of *PNKY* in human NSCs. Furthermore, we did not detect *Pnky* amplicons larger than the expected size in our *in cellulo* SHAPE-MaP studies, though it remains possible that isoforms similar to human *PNKY* exist in the mouse.

Typically, RNA structure-function studies begin by selecting certain folds or domains observed in the structural models for experiments that assess function. This approach benefits greatly from prior knowledge of the types of structures that are likely to possess function. However, for lncRNAs, there is currently very little information to guide the design of structure-function studies. More generally, it is not well-established that disrupting RNA structure will negatively impact the function of some (or most) lncRNAs. Therefore, to move beyond assumptions, the lncRNA field needs an example of RNA structure-function analysis that is strongly coherent with the known lncRNA function.

Instead of selecting a handful of structures identified in our models of *Pnky* structure for functional studies, we implemented a strategy that interrogated the entire *Pnky* transcript at approximately 20 nt intervals. For this, we used LNA ASOs, which due to their high Tm (which would out-compete RNA-RNA duplexes) have been found to be structure-disrupting in studies of other noncoding RNAs and viral genomes ^66–69^. Often, the biological assays used to explore lncRNA structure-function are relatively crude and not very coherent with biological function (if known at all) in the animal. Therefore, we sought to study the role of *Pnky* structure for the neuronal differentiation which despite being a complex cellular phenotype, is the most relevant. We reasoned that this assay coupled with the systematic, relatively unbiased approach of LNA targeted perturbations would provide the experimental scale and robustness to (1) establish a structural basis to *Pnky* function in neurogenesis and (2) identify specific RNA regions and/or structures that underlie the function of *Pnky* in NSCs.

Our strategy of using 38 LNAs to target the 825 nucleotides of *Pnky* revealed a few general experimental observations that may guide their use for similar studies. First, a small fraction of LNAs will likely exhibit off-target effects, as we found in our transfection of NSCs that genetically lack *Pnky*. Interestingly, the off-target effect in our studies did not produce the “positive” phenotype of increased neurogenesis. Instead, the off-target effect appeared to be toxic to the production of neurons. Second, although LNAs do not directly induce RNaseH-mediated degradation, a fraction of LNAs may nevertheless reduce levels of the targeted transcript. For this subset of LNAs, the potential role of the targeted RNA structure cannot be disentangled from the influence of decreased transcript levels. Third, some LNAs will not affect levels of the targeted RNA nor produce a cellular phenotype, and these can be useful as additional negative controls. We note here that for *Pnky*, the seven LNAs that did not produce a statistically significant effect upon neurogenesis in NSCs exhibited a trend of increased neuronal production, so we do not interpret the lack of phenotype as a lack of function of the targeted structure.

In our studies conducted in primary mouse NSC cultures, the positive neurogenic phenotype of 18 LNAs support the hypothesis that the structure of *Pnky* – independent of the level of the lncRNA transcript – plays a critical role in the regulation of neuronal production. Transfection of primary NSC cultures generally inhibits neuronal production, likely due to the cellular stresses of this experimental perturbation, and so the phenotype of increased neurogenesis with transfection of specific LNAs is noteworthy.

Our discovery that lncRNA *Pnky* structure underlies the biological regulation of neurogenesis represents a significant, foundational advance towards the broader understanding of lncRNA mechanisms. Mapping the location of the LNAs to both *in vitro* and *in vivo* conformations provided key insights that inform the next steps of experimental investigations into the structure-driven molecular mechanisms of *Pnky* and lncRNAs, in general. A subset of the LNAs mapped to locations with evidence of tertiary structure, including a predicted pseudoknot, which warrants further experimental interrogation to decipher their functional role. With the positive phenotypes of LNAs that target the modules that were similar between the *in cellulo* and *in vitro* models, these regions may be hypothesized to play a role in structure-function that is independent of other nuclear factors. LNAs mapping to the Modules 4 and 7 that exhibit distinct conformation *in cellulo* prioritize regions that may interact with nuclear proteins, such as PTBP1, which was previously shown to interact with *Pnky in vitro* and *in vivo* ^5^. Overall, this comprehensive approach to studying the structure of *Pnky* lays important next steps on the experimentally informed path to understanding lncRNA function and mechanism.

## RESOURCE AVAILABILITY

### Lead contact

Further information and requests for resources and reagents should be directed to and will be fulfilled by the lead contact, Daniel Lim.

### Materials availability

Plasmid, mouse lines and cell lines generated in this study will be made available upon request. We may require a payment and/or a completed materials transfer agreement in case there is potential for commercial application.

### Data and code availability

SHAPE-MaP and Terbium-seq data have been deposited in the NCBI SRA database under the identifier BioProject ID: PRJNA1263515.

This paper does not report original code.

Any additional information required to reanalyze the data reported in this paper is available from the lead contact upon request.

## ACKNOWLEDGEMENTS

The authors thank members of the Lim and Pyle lab for helpful discussions. We would like to thank Sarah Fergione and Olga Fedorova (Pyle Lab, Yale University) for their support with LNA ASO synthesis. We thank Sandra Chang (Lim Lab, UCSF) for the assistance with animal husbandry and administrative expertise for material transfer. This work was supported by Howard Hughes Medical Institute (A. M. P is an Investigator in the Howard Hughes Medical Institute), NIH grant T32GM007223-45 to S.P and NIH award 1R01NS124881 and Veterans Affairs 5I01 BX000252 to D.A.L.

## Author contributions

P.S and S.P designed, performed and analyzed the experiments. R.CA.T performed and analyzed SV-AUC and *in vitro* SHAPE experiments. H.C performed the cell-culture experiments and quantifications. R.E.A contributed to the design and generation of the plasmid constructs and transgenic mouse lines. L.T performed data analysis. D.A.L and A.M.P conceptualized and supervised the study and acquired funding. P.S. compiled figures and wrote the manuscript with D.A.L. All authors reviewed and edited the manuscript.

## METHODS DETAILS

### *In-vitro* transcription and purification

The *in vitro* transcription and semi-native purification of *Pnky* RNA was performed as previously described in ^25^. In brief, RNAs were transcribed by runoff transcription using T7 RNA polymerase (Ref) in a buffer containing 12mM MgCl_2_, 40mM Tris-Cl pH8, 2mM Spermidine, 10mM NaCl, 0.01% Triton X-100, 10mM DTT, 5μl SUPERase-In and 3.6mM of each NTP. The reactions were incubated at 37°C for 2 hours. Thereafter, 2U of TURBO DNase (2U/ul, Life Tech) was added and the mixture was incubated at 37°C for 30 min. To chelate excess divalent ions, 5μl of 0.1M EDTA was added. The RNA was buffer exchanged into filtration buffer (50mM MOPS-KOH pH 7, 150mM KCl) using a 100-kDa Amicon Ultra filtration column at room temperature. The RNA was purified using a self-packed 24ml Sephacryl S-400 gel filtration column equilibrated in filtration buffer. RNA from the peak fraction was folded by incubating 15mM MgCl_2_ at 37°C for 40 min.

### Sedimentation Velocity-Analytical Ultracentrifugation

SV-AUC experiments were performed with a Beckman XL-1 centrifuge and an AN-50 Ti Analytical Rotor. For each condition, 500μl of the RNA at an A260 of 0.6 was used. RNA was folded as described above. For each condition, a 500μl of buffer was also prepared. AUC cells were cleaned thoroughly with RNase Zap and 100% ETOH, dried, and assembled according to the manufacturer’s instructions. Using a round gel loading tip, 2x 210μl of buffer was added to the left side of the cell chamber. Similarly, 2x 195μl of RNA was added to the right side of the cell chamber. Cells were sealed according to the manufacturer’s instructions, placed into the rotor, and equilibrated to 20°C for 1.5-2 hours under vacuum. Thereafter, cells were spun at 25,000 rpm for 16 hours for a total of 120 scans. Data was analyzed in Sedfit ^70^ using the continuous distribution model Hydrodynamic radii (R_h_) were calculated in Sedfit using a partial specific volume of 0.53 cm^3^/g and a hydration of 0.59 g/g. Buffer viscosity and density were estimated using Sednterp and figures were made using Gussi c(s)

### In vitro SHAPE

For probing, 2μg of RNA was incubated with 15mM MgCl_2_ in a final volume of 20μl and incubated at 37°C for 40 min. Thereafter, 18μl of folded RNA was transferred to a tube containing either 2μl of 100mM 1M7 (final concentration 10mM) or pure DMSO as a control and incubated at 37°C for 10 min followed by quenching with 2μl of 1M DTT. All RNA samples were purified using a Zymo RNA clean and concentrator kit according to manufacturer’s protocol and eluted in 15μl of nuclease-free water.

### Animals

All animal related research protocols comply with the relevant ethical regulations approved by the University of California, San Francisco Institutional Animal Care and Use Committee. C57BL/6J mice used in this study were maintained in the University of California, San Francisco Laboratory Animal Resource Center under approved protocol (AN195649-01C). Mice were group-housed in cages with sterile bedding and *ad libitum* food and water. Standard conditions of 12-h dark/light cycle, humidity (30-70%) and temperature (20-26°C) were maintained. Mice of both sexes were used for all experiments and were used for derivation of primary cell culture at postnatal day 7. Mouse strains used in this study are *Pnky* ^+/+^ (WT), *Pnky*^-/-^(knockout) and *Pnky*^+/+^; BAC-*Pnky* (overexpression). All animals were genotyped by PCR for *Pnky*^+^, and *Pnky*^-^ alleles using the following primers: *Pnky* GT Forward and Reverse 1 and 2 primers. Reaction products: 120 bp (*Pnky*-KO), 221 bp (*Pnky*-WT). Since BAC-*Pnky* contains unaltered *Pnky*, this will also produce the 221-bp product. Primers for BAC-*Pnky* produce a 416 bp reaction product and no amplification in the absence of BAC-*Pnky*.

### Ventricular subventricular zone (V-SVZ) neural stem cell culture

The brains of P7 mice were dissected out of the skull, and then a coronal slab of approximately 0.5 mm in thickness was obtained manually. Dissections were performed in ice-cold L15 medium (ThermoFisher 11415064) to collect the V-SVZ region along the lateral walls of the lateral ventricles. The tissue was dissociated with 300 mL of 0.25% trypsin-EDTA (ThermoFisher 25200056) with occasional agitation for 20 min at 37 °C, and then the trypsin was quenched with 600 mL of N5 growth medium (see below). Cells were mechanically dissociated by trituration and then pelleted by centrifugation at 300 g for 5 min. Cells were resuspended in fresh N5 growth medium and plated in one well of a 12-well tissue culture plate per mouse. Cells from mice of same genotype from each litter (both sexes included) were pooled together. Cells were grown in N5 growth medium: DMEM/F-12 with GlutaMAX (ThermoFisher 10565018) supplemented with 5% fetal bovine serum (Fisher Scientific SH30070.03), N2 supplement (ThermoFisher 17502048, 1:100), 35 mg/mL bovine pituitary extract (ThermoFisher 13028014), antibiotic-antimycotic (ThermoFisher 15240062, 1:100), 20 ng/mL epidermal growth factor (EGF, PeproTech AF-100-15), and 20 ng/mL basic fibroblast growth factor (bFGF, PeproTech 100–18 B). For routine passaging, cells were grown to hyper confluence and then dissociated using 0.25% trypsin-EDTA (ThermoFisher 25200056) for 3–5 minutes in the incubator. This was diluted with an equal volume of growth medium, and the cells were dissociated by trituration. Cells were pelleted as described above, then resuspended in fresh culture medium to plate in 2 times the original culture area. For differentiation, at passage 4-5 cultures were grown to hyperconfluence and then the medium was replaced with the differentiation media (N5 media without EGF or FGF and 2% serum) for 5 days. All cells were grown in humidified incubators at 37°C in 5% CO2.

### In-cell SHAPE

SVZ NSCs (*Pnky*^+/+^; BAC-*Pnky*) were grown in N5 growth medium were used. Approximately 70-90 million cells were used for each probing condition. For all probing conditions, media was aspirated, cells were washed once with DPBS,and dislodged using Accutase. The cells were collected and centrifuged at 300g x 5min at 25°C. The supernatant was removed and to every ∼10 million cells 900μl of DPBS was added. Cells were resuspended by pipetting up and down to ensure a homogenous cell suspension. 100ul of freshly dissolved 1M7 (in DMSO) to a final concertation of 50mM or an equivalent volume of only DMSO (as control) was added to the cells. The reaction mixture was incubated at 37°C for 5 minutes and pelleted at 300g x 5min at 25°C. The resulting cell pellet was resuspended in nuclear fractionation buffer (10mM Tris-HCl, 140mM NaCl,1.5mM MgCl_2_, 0.5% (w/v) Igepal and 10mM Ribonucleoside vanadyl complex) and incubated on ice for 5-10 minutes. The mixture was centrifuged 1000g x 5min at 4°C. The supernatant (cytoplasmic fraction) was discarded, and the procedure was repeated two more times. In the final addition of the fractionation buffer, 0.5% (w/v) sodium-deoxycholate was added. The nuclear pellet was resuspended in Trizol and extracted using Direct-zol RNA Purification Kits (Zymo Research) according to manufacturer’s protocol. The RNA was eluted in ME buffer (8mM MOPS pH 6.5, 1mM EDTA pH 8.5). Nuclear RNA was ribosome depleted using ThermoFisher (K155001) Ribominus kit according to manufacturer’s protocol.

### MaP Reverse Transcription

For each probing condition, ∼2000ng of IVT RNA or ∼800-1000ng of nuclear enriched, ribodepleted RNA was mixed with 2 pmol of RT primer in a final volume of 10μl and annealed at 90°C for 1 min followed by 30°C for 2 min. Thereafter, 20U/μl of Superscript II, 2U/μl of SUPERase-In RNase inhibitor, and SHAPE-Map Buffer (50mM 1M Tris-HCl pH 7.5, 75mM KCl, 10mM DTT, 0.5mM dNTPs, 6mM Mn^2+^) were added and the RT reactions were incubated at 42°C for 3 hours. The resulting cDNA was purified using a 1.8:1 ratio of AMPure XP beads to sample according to manufacturer’s protocol. Purified cDNA was eluted in 12μl of nuclease-free water

### PCR and amplicon generation

Next, amplicons were generated by using 0.5mM of gene-specific forward and reverse PCR primers (Appendix, Table 1), 1x Q5 reaction buffer, 0.2mM dNTPs, and 0.02U/ul Q5 HF Hot start DNA Polymerase and 10μl of purified cDNA in a final volume of 50uμ. Touch down PCR cycles were used to enhance specificity with a final annealing temperature of 63°C for Amplicon 1 or 60°C for Amplicon 2. PCR samples were purified using 1:1 AMPure XP beads to sample ratio and eluted in 12μl of nuclease-free water.

### SHAPE sequencing library prep

Amplicon concentrations were determined using a Qubit dsDNA kit according to manufacturer protocol and diluted to 0.2ng/ul. To generate sequencing libraries the NexteraXT DNA library prep kit was used with 1/5 of the manufacturer’s recommended volumes. Libraries were once more cleaned using a 1:1 AMPure beads to sample ratio.

### Sequencing

Library concentrations were determined using a Qubit dsDNA HS Assay Kit and a BioAnalyzer High Sensitivity DNA Analysis. Libraries were diluted, pooled and sequenced using a NextSeq 500/550 or NextSeq 2000 platform.

### SHAPE-MaP data analysis

The sequencing data was analyzed using ShapeMapper2 ^50^, aligning reads to mouse *Pnky* transcript. SHAPE reactivities were computed by the ShapeMapper2 script by calculating mutation rate differences between SHAPE-modified and unmodified samples for each nucleotide of mouse *Pnky* and then normalizing the values into a final ∼0-2 reactivity scale.

### Secondary structure prediction

ShapeKnots ^46^ was first used in 600-nt windows to scan *Pnky* transcript for the presence of pseudoknots using *in vitro* derived SHAPE reactivities, in addition to a prediction using the full-length context. A pseudoknot was judged plausible if both strands of pseudoknotted helix had low (<0.4) SHAPE reactivities. A plausible pseudoknot was then used as constraints for the final secondary structure computation. A consensus secondary structure of the full-length mouse lncRNA *Pnky* was obtained using SuperFold 1.0 ^20^ with default settings along with *in vitro* SHAPE reactivities and the identified pseudoknot constraint. This consensus secondary structure map was used for comparison between *in vivo* and *in vitro* SHAPE results by mapping the SHAPE reactivity values from each experiment to every nucleotide position of the region of interest.

### Terbium probing

Terbium probing was performed by incubating 18μl of the folded RNA with 2μl of 5mM TbCl_3_ stocks prepared in monovalent buffers (final concentration 0.5mM) or 2μl of monovalent buffer (negative control) for 10 min at 25 °C. All reactions were quenched with the addition of 3μl of 50mM EDTA pH 8 and cleaned using a Zymo RNA clean and concentrator.

### Terbium library preparation

For each probing condition, the RNA was mixed with 0.5pmol of gene-specific primers (Appendix, Table 1) and brought to a volume of 7.5μl. To anneal primers, the mixture was heated at 90°C for 1 min followed by 30°C for 2 min. To initiate reverse transcription, 0.5ul of Marathon RT ^71^, 10μl of 2x Marathon Ammonium Sulfate buffer (100 mM Tris-HCl pH 8.3, 400 mM (NH_4_)_2_SO_4_, 4 mM MgCl2, 10 mM DTT and 40% glycerol), 1μl of 10mM dNTP mix (NEB), and 1ul of 10mM DTT were added and incubated at 42°C for 15 min. RNA was degraded with the addition of 1μl of 3M KOH, heated to 95°C for 5 min and snap cooled to 4°C for 5 min. Thereafter, 1μl of 3M HCl was added to neutralize the reaction. cDNA products from RT were purified using AMPure XP beads by adding 1.2x bead to sample ratio and incubating at room temp for 10 min. The beads were captured using a magnetic rack for 5 min and washed 3 times with 180μl of fresh 80% ETOH. The beads were air dried for 5 min and resuspended in 12μl of water to elute the cDNA. Thereafter, 3’ adaptor ligation was performed by mixing 8μl of purified cDNA with 0.2μl of 50μM 3’ adaptor (Appendix, Table 1), 1μl of 10mM ATP, 2μl of T4 RNA Ligase buffer, 8μl of 50% PEG 8000. The mixture was incubated at 25°C for 16 hours, followed by enzyme deactivation at 65°C for 15 min. Ligated products were purified with AMPure XP beads using a 1.2x bead to sample ratio. The products were PCR amplified 9 cycles with Q5 HF DNA polymerase using Illumina TruSeq forward primer and indexed reverse primers, with cycle times of 98°C for 10 sec, 62°C for 45 sec, and 72°C for 60 sec. PCR products were purified with 1.2x volume of AMPure XP beads.

### Terbium data analysis

All FASTQ files were processed using Cutadapt to remove Illumina adaptor sequences and aligned to the respective RNA sequence using HISAT2 (v2.10). Stop information was extracted using the RTEventsCounter.py script ^72^. The probability of stop per nucleotide was calculated as the number of stops divided by the sum of the total number of read-through events plus the number of stops. Probabilities were background subtracted against a no probe control. The data was normalized and scaled as in ^24^.

### LNA and Gapmer ASO design

LNAs were designed with three consecutive LNA bases at the 5’ and 3’ ends with stretches of upto three consecutive unlocked bases. The design was based on thermodynamic properties such as length, % GC content and LNA:RNA duplex T_m_. All LNA oligos were synthesized on a MerMade 12 RNA-DNA synthesizer (BioAutomation) using DNA and LNA phosphoramidites (TxBio) and Glen UnySupport 1000 (GlenResearch). The synthesis cycle was modified to increase oxidation step to 3 min as recommended by the manufacturer. Removal of oligonucleotides from the polymer support and base deprotection were carried out in 28 to 30% aqueous ammonium hydroxide: 40% aqueous methylamine (1:1) at 65 °C for 2 h. Then, the resulting oligo solutions were dried using Speedvac, redissolved in 750 μL of sterile deionized water, and desalted using GlenPak 1.0 columns (Glen Research). Scrambled LNA was designed for control. Gapmer ASOs (negative control and against *Pnky* RNA) was design using the Qiagen Antisense LNA Gapmer design tool and procured from Qiagen.

### LNA and Gapmer ASO transfection

SVZ NSCs (at passage 4-5) were plated in poly-D-lysine and laminin coated 48-well culture plates (20-25,000 cells per well) for RNA extraction or 96 well Mattek (P96GC-1.5-5-F, No. 1.5 Coverslip | 5 mm Glass Diameter) glass bottom plate (10-12,000 cells per well) for differentiation and imaging in N5 proliferation medium, one day before transfection. Transfection was performed in 300μl total volume for 48-well plates and 150μl total volume for 96 well plates such that the final concentration of the LNA ASO is 100nM and Gapmer ASO is 50nM. Appropriate volume of LNA or Gapmer ASO was diluted in Opti-MEM reduced serum media from 20μM stock. Oligofectamine reagent (Invitrogen Cat 12252011) was diluted in the Opti-MEM media and incubated for 5-10 min. The diluted oligonucleotides were combined with diluted Oligofectamine, mixed and incubated for 15-20 min at room temperature. While complexes are forming, the growth media is removed and replaced with 100ul and 50ul of N5 media without serum respectively. The transfection reagent with the LNAs/ASOs are added to the cells and incubated for 4 hrs at 37°C in the CO_2_ incubator. Thereafter, N5 media with 3X the normal concentration of the serum is added without removing the transfection mixture.

### NSC differentiation assay

21-24 hrs post transfection with LNA or ASO, the N5 media with the oligonucleotides is removed and the differentiation media N6 (without the growth factors) is added to each well. Cells are allowed to undergo differentiation for 5 days.

### Immunocytochemistry (ICC)

After 5 days of differentiation, the cells are washed 1X with PBS and fixed with 4% paraformaldehyde for 30 min at RT. Post fixation, cells are washed 3X with PBS. citrate antigen retrieval was performed using heated 10 mM sodium citrate (pH 6.0) for 10-15 min in a slide box. The sodium citrate was then dumped off and the slides were rinsed in PBS. ICC were performed using blocking buffer consisting of PBS with 1% BSA (Millipore Sigma), 0.3-M glycine (Thermo Fisher Scientific), 0.3% TritonX-100 (Millipore Sigma), and either 10% normal goat serum or 10% normal donkey serum (Jackson ImmunoResearch Laboratories). Cells were incubated in blocking buffer for 1 hr at RT, followed by the incubation in primary antibody Tuj1 (Biolegend, #Cat 801201) diluted (1:1000) in blocking buffer for 2 hrs at RT. Slides are washed 3X in PBS at RT for 5 min, followed by the incubation in secondary antibodies (Alexa Fluor antibodies from Thermo Fisher Scientific, 1:500) and DAPI (Thermo Fisher Scientific, 1:500) diluted in blocking buffer for 45 min at RT. This is followed by 3X wash in PBS and then coverslipped using the AquaMount.

### Microscopy imaging and quantification

Cells were imaged using Leica Epifluorescence microscope. For each transfection condition, experiments were performed in independent triplicates. For each well, 3-4 non-overlapping fields of view were imaged. All images were processed using Fiji and quantified using the Imaris software (spots function where the diameter of the spot was set to 8.00) by an experimenter blinded to the target location of LNA probes on *Pnky* RNA. Total number of DAPI and Tuj1 positive spots were counted to obtain % Tuj1+ normalized to total DAPI positive nuclei.

### RNA isolation

21-24 hrs post transfection with LNA or ASO, total RNA was isolated from cultured cells using Trizol (Thermo Fisher Scientific) and purified using Direct-zol RNA Purification Kits (Zymo Research). DNase digestion was performed as suggested.

### RT-qPCR

cDNA synthesis was performed using the Transcriptor first Strand cDNA synthesis kit (Roche). qRT-PCR was performed using SYBR Green (Roche) on a LightCycler 480 II (Roche). Relative gene expression was calculated using the ^ΔΔ^Ct method, using GAPDH as a housekeeping gene for differential gene expression analyses. All qRT-PCR assays were performed using technical triplicate wells.

### Figure Preparation

Graphs were generated using PRISM (GraphPad). Figures were generated using Adobe Illustrator and schematics using BioRender.

