## Supplementary Information for "Structural basis for the function of long noncoding RNA *Pnky* in neural stem cells"

**Document S1. Figures S1–S6 and Tables S1-S4**

### SUPPLEMENTAL FIGURE LEGENDS

**Figure S1. Amplicons for *in vitro* SHAPE MaP, related to Figure.2**

(A) Schematic for amplicon PCR primer design – two sets of overlapping primers were designed to amplify the fully spliced *Pnky* RNA (825bp).

(B) Gel image showing the products of the amplicon PCR for the two replicates of DMSO and 1M7 treated samples.

**Figure S2. Quality metrics for the *in vitro* SHAPE MaP datasets, related to Figure. 2**

(A) Comparison of mutation rates of 1M7 modified and unmodified samples for n= 2 independent replicates. Mean +/- SD, ****p <0.0001; unpaired two-tailed Student's t test.

(B) Correlation plot of normalized SHAPE reactivities from 2 replicates. Linear regression fit to the data is represented by the line. Pearson's correlation for the dataset is indicated.

(C) SHAPE reactivity profile for 1M7 treated replicates of *in vitro* transcribed *Pnky* RNA.

(D) Strandedness of low, moderate and highly SHAPE reactive nucleotides in the two replicates of *in vitro* SHAPE datasets.

(E) Shannon entropy is significantly lower in the model with the pseudoknot included ***p <0.001; unpaired two-tailed Student's t test.

See also Table S1.

**Figure S3. Quality metrics for the terbium sequencing datasets, related to Figure. 3**

(A) Bar plot showing reactivity profile obtained when probing *in vitro* transcribed and folded *Pnky* RNA for the two independent replicates.

(B) Correlation plot of normalized terbium reactivities from 2 replicates. Linear regression fit to the data is represented by the line. Pearson's correlation for the dataset is indicated.

(C) Secondary structure of the *Pnky RNA* displaying sites of strong $Tb^{3+}$cleavage (red) for the two independent replicates.

See also Table S1.

**Figure S4. Quality metrics for the *in-cell* SHAPE MaP datasets, related to Figure. 4**

A) Gel image showing the products of the amplicon PCR for the two independent biological replicates of DMSO and 1M7 treated samples.

(B) Comparison of mutation rates of 1M7 modified and unmodified samples for two independent replicates. Mean +/- SD, $^{**}p < 0.01$, $^{***}p < 0.001$; unpaired two-tailed Student's t test

(C) showing the SHAPE reactivity profile for 1M7 treated replicates for the two independent *in-cell* SHAPE experiments.

(D) Correlation plot of normalized SHAPE reactivities from two biological replicates from neural stem cells. Line represents the linear regressions fit to the data. Pearson's correlation for the datasets is shown.

(E) Stranded-ness of low, moderate and highly SHAPE reactive nucleotides in the two biological replicates of *in-cell* SHAPE experiments.

See also Table S1.

**Figure S5. LNA and Gapmer ASO transfection in *Pnky*-WT NSCs and differentiation assay, related to Figure.5**

(A) Schematic of the LNA and Gapmer ASO transfection in the NSCs and differentiation assay**.**

(B) Representative images of the Tuj1 ICC with DAPI nuclear stain in day5 differentiated *Pnky*-WT V-SVZ neural stem cell primary culture for different ASO/LNA transfections. Scale bar 50μm.

See also Table S3.

**Figure S6. LNA ASO transfection in *Pnky*-KO NSCs and differentiation assay, related to Figure.5**

(A) Schematic of the LNA ASO transfection in the NSCs and differentiation assay**.**

(B) Representative images of the Tuj1 ICC with DAPI nuclear stain in day5 differentiated *Pnky*-KO V-SVZ neural stem cell primary culture for different ASO/LNA transfections. Scale bar 50μm.
See also Table S3.

### SUPPLEMENTAL TABLES

**Table S1.** Quality metrics of chemical probing sequencing data (SHAPE MaP) related to Figures 2-4.
**Table S2.** Module locations/nucleotides (*in vitro* vs *in-cell*), related to Figures 2 and 4.
**Table S3.** List of LNAs used: sequence, targeting nucleotide positions, Tm. Related to Figures 5 and 6.
**Table S4.** List of primers- genotyping, amplicon primers, RT primers, qPCR primers

Figure S1

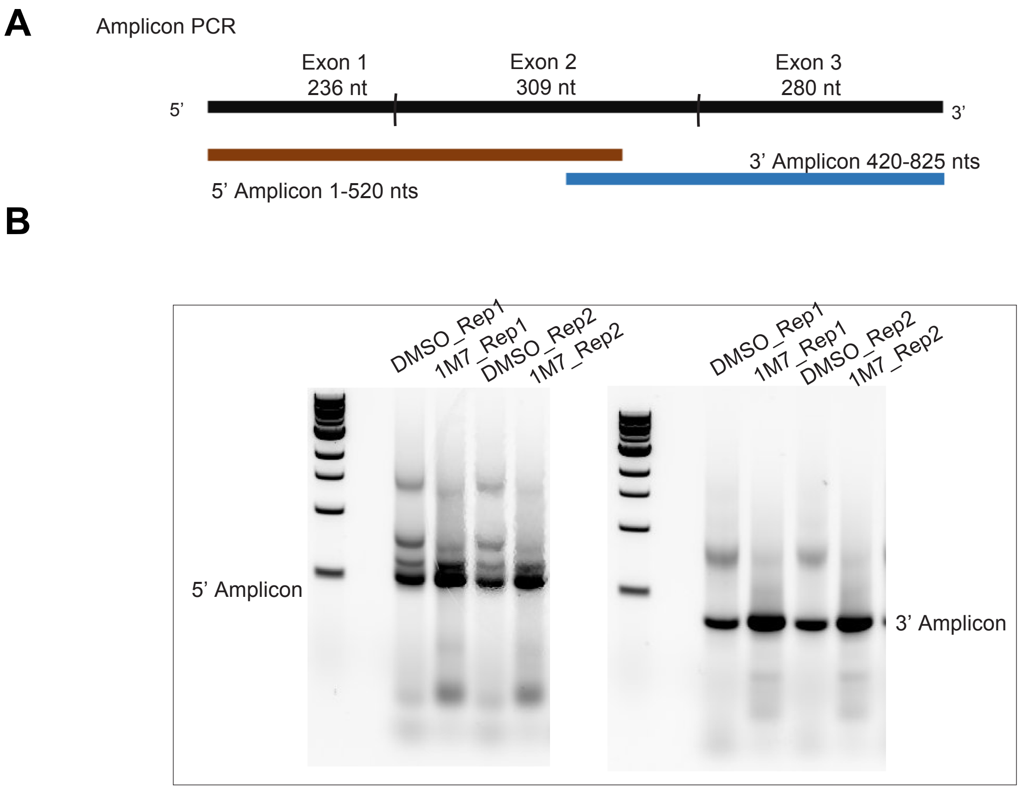

**Figure S1:Amplicons for the *in vitro* SHAPE MaP related to Figure.2**

(A) Schematic for amplicon PCR primer design – two sets of overlapping primers were designed to amplify the fully spliced *Pnky* RNA (825bp).

(B) Gel image showing the products of the amplicon PCR for the two replicates of DMSO and 1M7 treated samples.

Figure S2

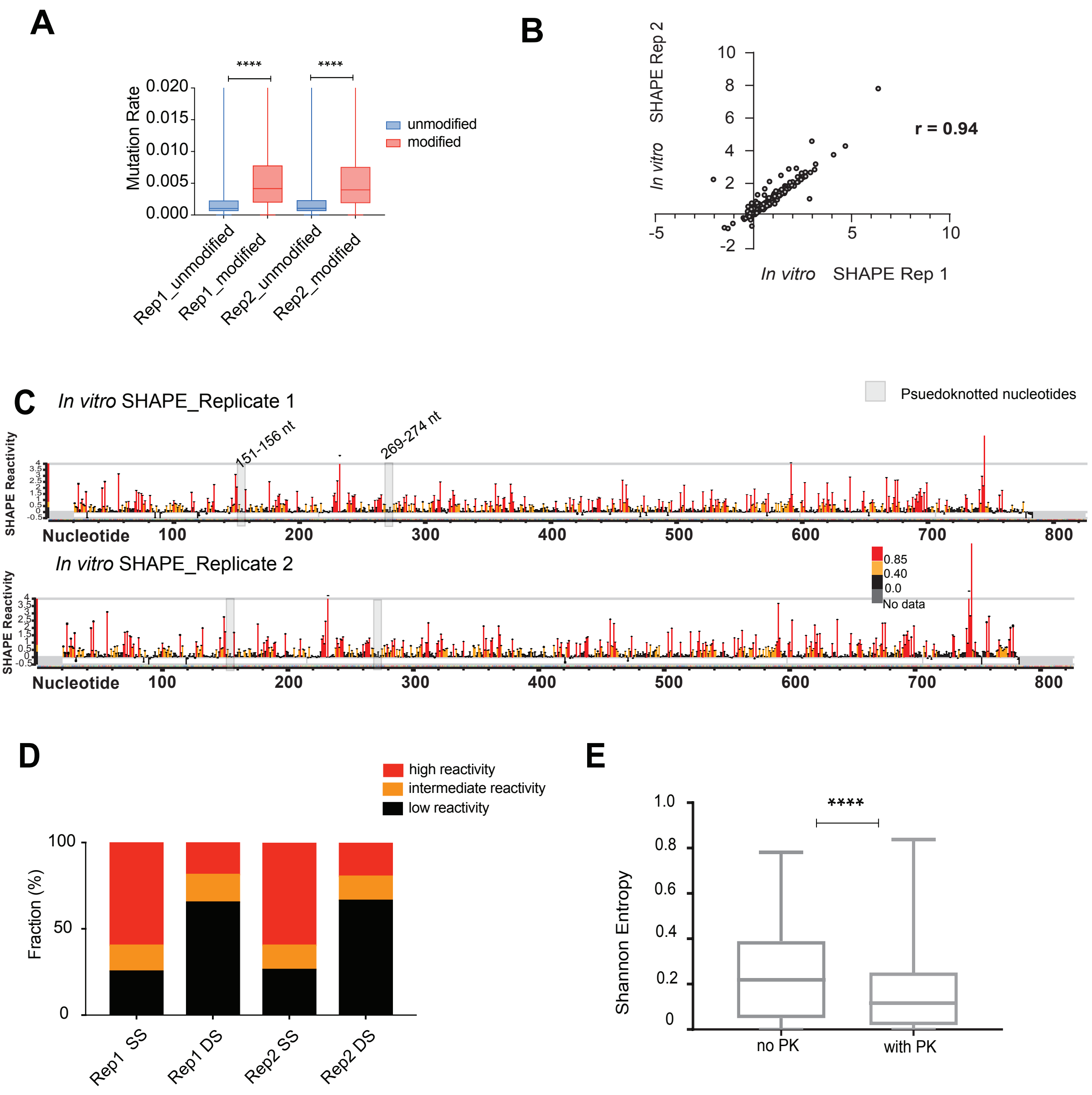

**Figure S2: Quality metrics of *in vitro* SHAPE MaP datasets, related to Figure. 2.**
(A) Comparison of mutation rates of 1M7 modified and unmodified samples for n= 2 independent replicates. Mean +/- SD, *****p* <0.0001; unpaired two-tailed Student's t test.
(B) Correlation plot of normalized SHAPE reactivities from 2 replicates. Linear regression fit to the data is represented by the line. Pearson's correlation for the dataset is indicated.
(C) SHAPE reactivity profile for 1M7 treated replicates of *in vitro* transcribed *Pnky* RNA.
(D) Strandedness of low, moderate and highly SHAPE reactive nucleotides in the two replicates of *in vitro* SHAPE datasets.
(E) Shannon entropy is significantly lower in the model with the pseudoknot included *****p* <0.0001; paired two-tailed Student's t test.
See also Table S1.

Figure S3

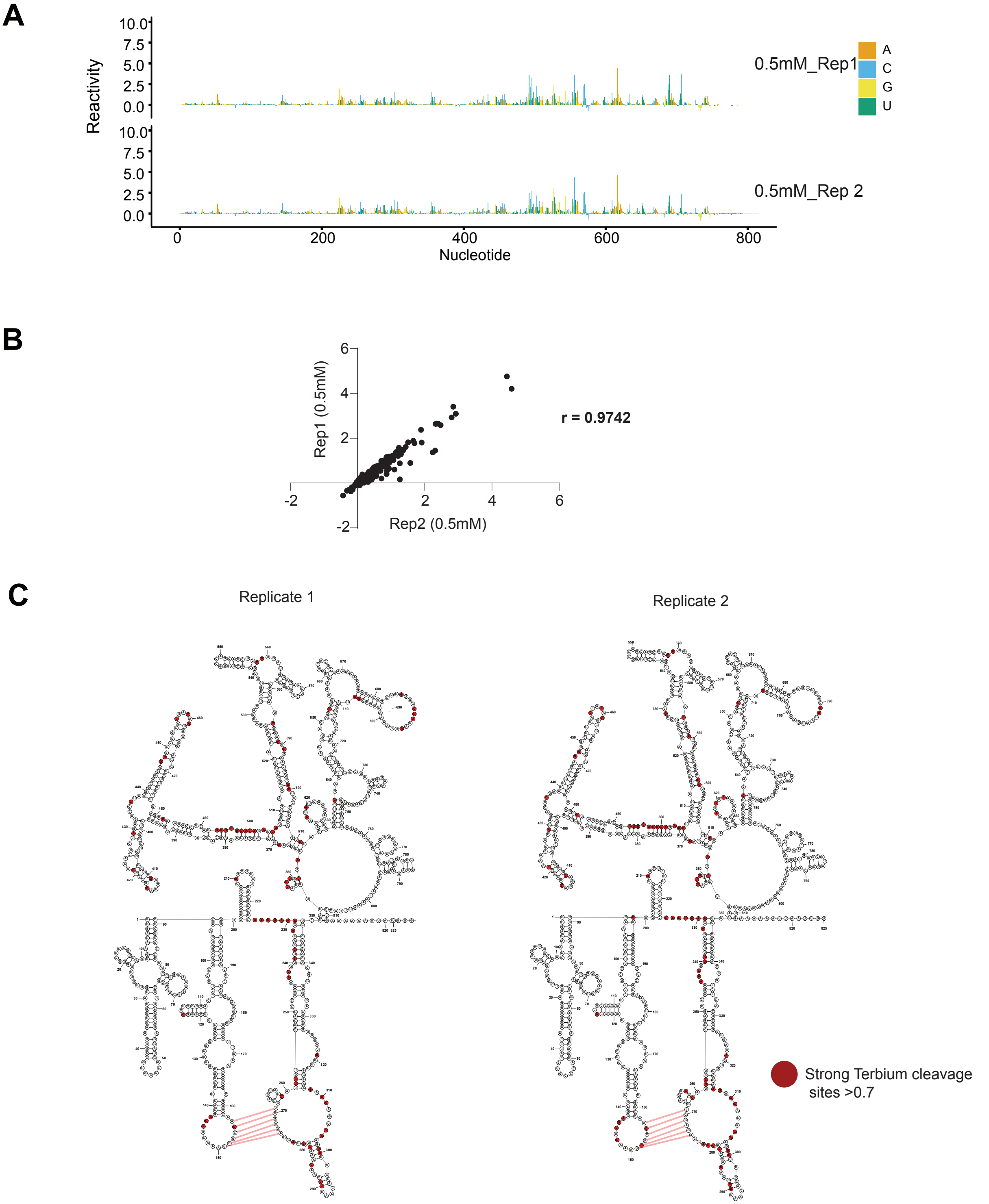

**Figure S3.Quality metrics for the terbium sequencing datasets, related to Figure 3.**
(A) Bar plot showing reactivity profile obtained when probing in vitro transcribed and folded *Pnky* RNA for the two independent replicates.
(B) Correlation plot of normalized terbium reactivities from 2 replicates. Linear regression fit to the data is represented by the line. Pearson's correlation for the dataset is indicated.
(C) Secondary structure of the *Pnky* RNA displaying sites of strong $Tb^{+3}$cleavage (red) for the two independent replicates.
See also Table S1.

Figure S4

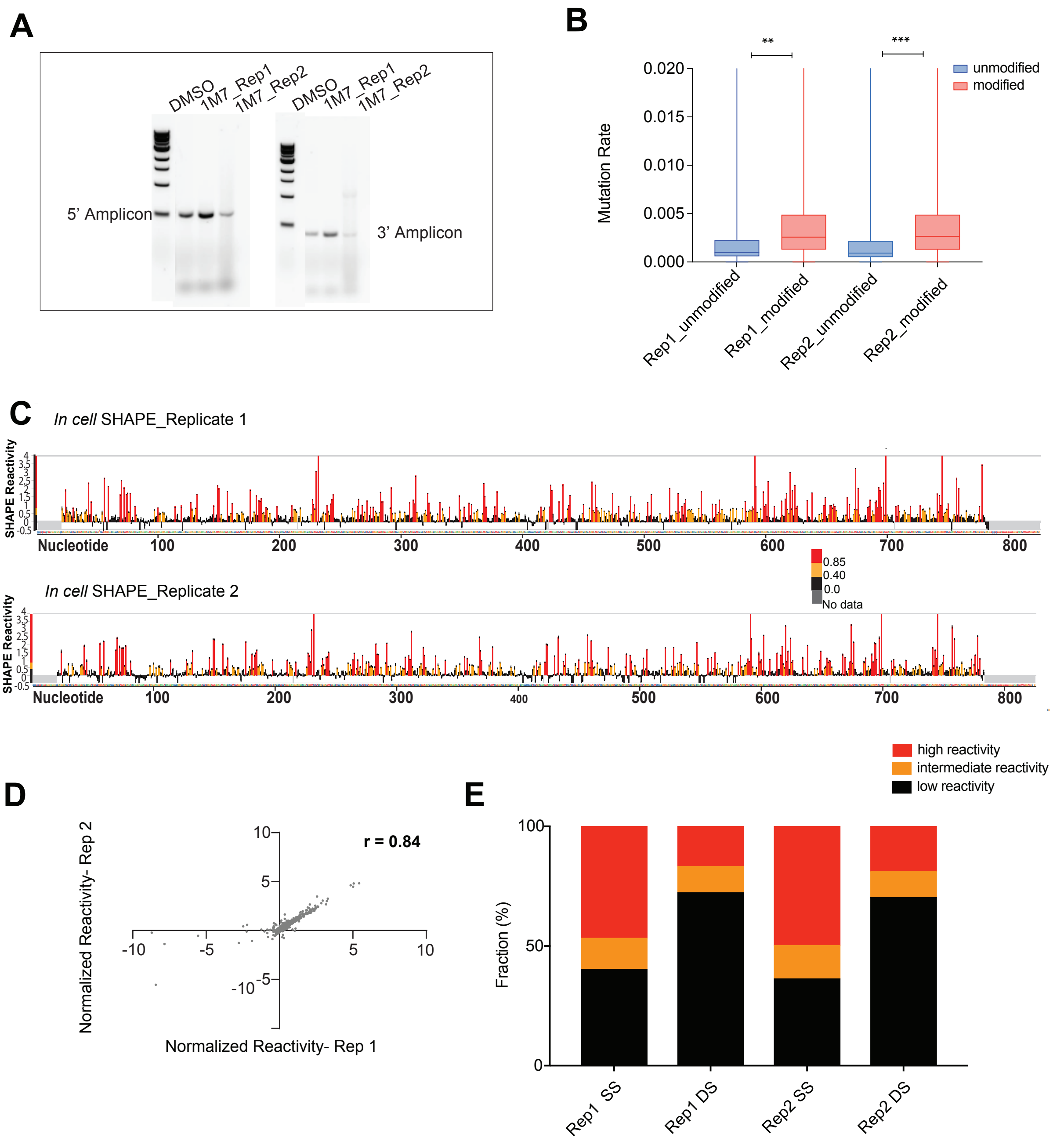

**Figure S4. Quality metrics for the *in-cell* SHAPE datasets, related to Figure 4.**
(A) Gel image showing the products of the amplicon PCR for the two independent biological replicates of DMSO and 1M7 treated samples.
(B) Comparison of mutation rates of 1M7 modified and unmodified samples for two independent replicates. Mean +/- SD, $^{**}p < 0.01$, $^{***}p < 0.001$; unpaired two-tailed Student's t test.
(C) showing the SHAPE reactivity profile for 1M7 treated replicates for the two independent *in-cell*SHAPE experiments.
(D) Correlation plot of normalized SHAPE reactivities from two biological replicates from neural stem cells. Line represents the linear regressions fit to the data. Pearson's correlation for thedatasets is shown.
(E) Stranded-ness of low, moderate and highly SHAPE reactive nucleotides in the two biological replicates of *in-cell* SHAPE experiments.
See also Table S1.

Figure S5

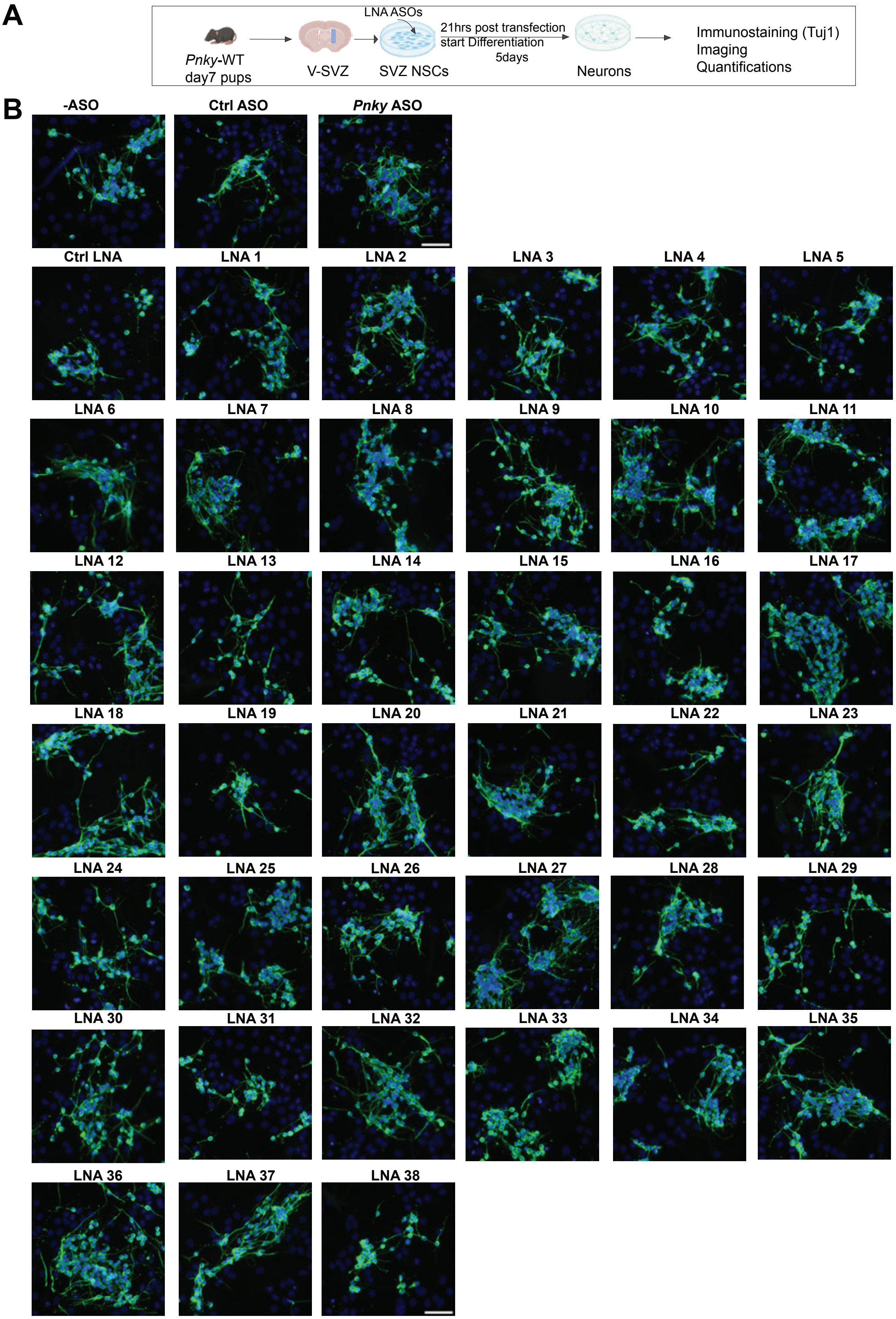
A
LNA ASOs
21hrs post transfection
start Differentiation
5days
Pnky-WT
day7 pups
V-SVZ
SVZ NSCs
Neurons
Immunostaining (Tuj1)
Imaging
Quantifications
B
-ASO
Ctrl ASO
Pnky ASO
Ctrl LNA
LNA 1
LNA 2
LNA 3
LNA 4
LNA 5
LNA 6
LNA 7
LNA 8
LNA 9
LNA 10
LNA 11
LNA 12
LNA 13
LNA 14
LNA 15
LNA 16
LNA 17
LNA 18
LNA 19
LNA 20
LNA 21
LNA 22
LNA 23
LNA 24
LNA 25
LNA 26
LNA 27
LNA 28
LNA 29
LNA 30
LNA 31
LNA 32
LNA 33
LNA 34
LNA 35
LNA 36
LNA 37
LNA 38

**Figure S5: LNA and gapmer ASO transfection in the *Pnky*-WT NSCs and differentiation assay. Related to Figure 5.**

(A) Schematic of the LNA and Gapmer ASO transfection in the NSCs and differentiation assay.

(B) Representative images of the Tuj1 ICC with DAPI nuclear stain in day5 differentiated *Pnky*-WT V-SVZ neural stem cell primary culture for different ASO/LNA transfections. Scale bar 50um.

See also Table S3.

Figure S6

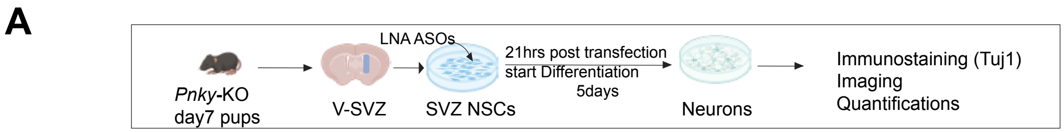
A
Pnky-KO
day7 pups
V-SVZ
LNA ASOs
SVZ NSCs
21hrs post transfection
start Differentiation
5days
Neurons
Immunostaining (Tuj1)
Imaging
Quantifications

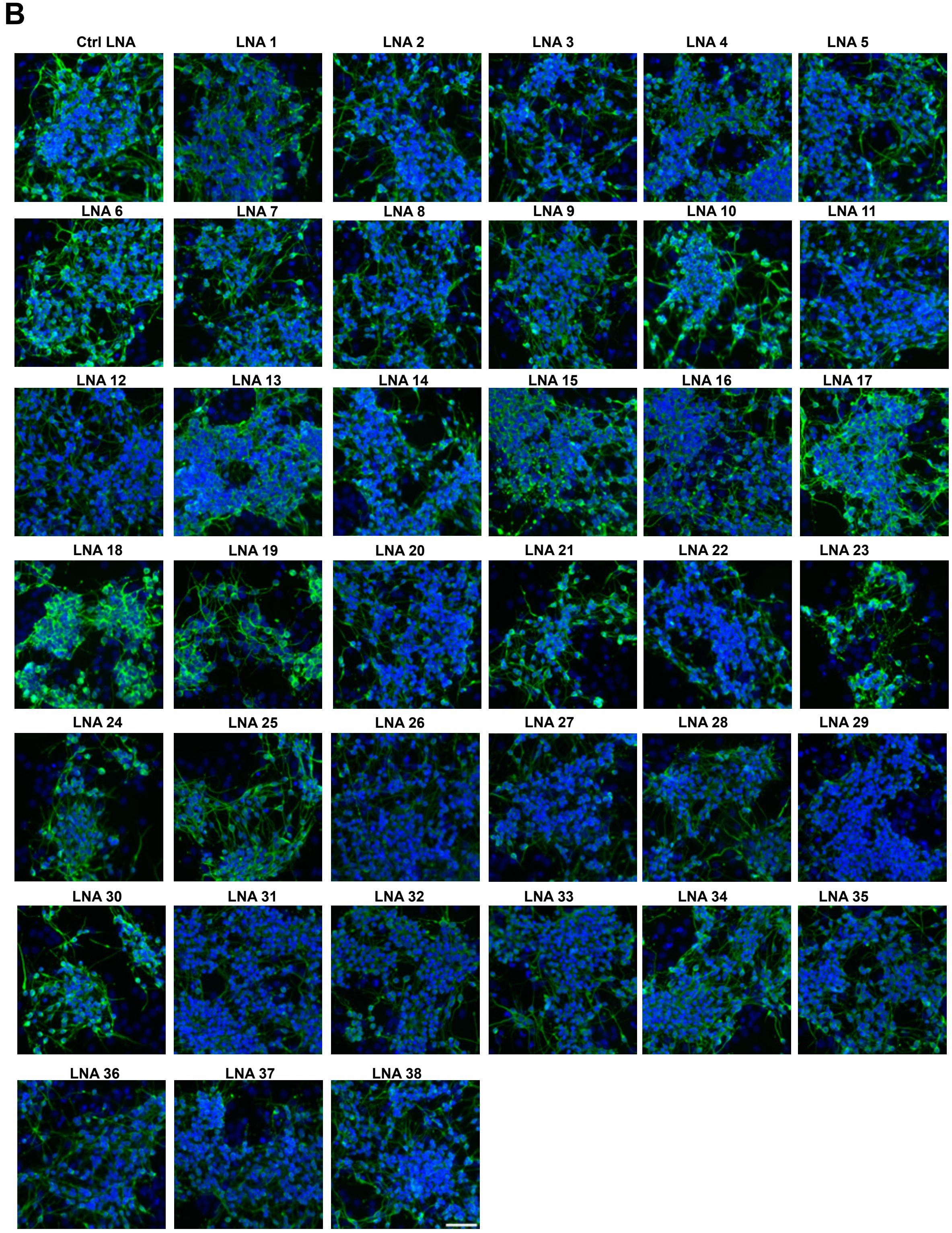
B
Ctrl LNA
LNA 1
LNA 2
LNA 3
LNA 4
LNA 5
LNA 6
LNA 7
LNA 8
LNA 9
LNA 10
LNA 11
LNA 12
LNA 13
LNA 14
LNA 15
LNA 16
LNA 17
LNA 18
LNA 19
LNA 20
LNA 21
LNA 22
LNA 23
LNA 24
LNA 25
LNA 26
LNA 27
LNA 28
LNA 29
LNA 30
LNA 31
LNA 32
LNA 33
LNA 34
LNA 35
LNA 36
LNA 37
LNA 38

**Figure S6. LNA ASO transfection in the *Pnky*-KO NSCs and differentiation assay. Related to Figure 5.**

(A) Schematic of the LNA ASO transfection in the NSCs and differentiation assay.

(B) Representative images of the Tuj1 ICC with DAPI nuclear stain in day5 differentiated *Pnky*-KO V-SVZ neural stem cell primary culture for different ASO/LNA transfections. Scale bar 50um.

See also Table S3.

**Table S1. Quality metrics of chemical probing sequencing data (SHAPE seq), related to Figures 2-4.**

| Libraries | Read depth check (%) | Mutation rate check (%) | High background check % | No. of highly reactive nucleotides check (%) |
|---|---|---|---|---|
| **SHAPE Mapper threshold** | >80% | >50% | <5% | >8% |
| **In cell Rep 1** | 97.8% | 85.1% | 0.4% | 6.7% |
| **In cell Rep 2** | 97.8% | 90.8% | 0.4% | 8% |
| **In vitro Rep 1** | 97.8%` | 94.6% | 0.4% | 20.3% |
| **In vitro Rep 2** | 97.8% | 93.8% | 0.4% | 19.1% |

**Table S2. Module locations/nucleotides (*in-cell* vs *in vitro*), related to Figures 2 and 4.**

| Modules | *In vitro* model | *In-cell* model |
|---|---|---|
| 1 | 1-91 | 1-91 |
| 2 | 92-200 | 92-200 |
| 3 | 201-224 | 201-224 |
| 4 | 232-350 | 232-350 |
| 5 | 371-505 | 371-505 |
| 6 | 507-608 | 507-608 |
| 7 | 613-774 | 609-783 |

**Table S3. List of LNAs used: targeting nucleotide positions, sequence, Tm. Related to Figures 5 and 6.**

| LNA | Nucleotide position | Sequence | length | Tm |
|---|---|---|---|---|
| **Ctrl LNA** | | 5'+G+T+GTA+ACA+CGT+CTA+TAC+GC+C+C+A 3' | 22 | 81 |
| **1** | 1-22 | 5' +A+C+CAG+AGG+AAG+TTG+CTT+C+T+C 3' | 20 | 78 |
| **2** | 23-42 | 5' +G+T+AGG+TTA+CAC+CTC+CAG+A+A+G 3' | 20 | 76 |
| **3** | 43-62 | 5' +C+A+GTA+TAT+CCT+ACT+GGG+C+A+C 3' | 20 | 78 |
| **4** | 63-83 | 5' +C+G+TGG+ACA+TTT+CAC+AA+CC+C+G+G 3' | 21 | 86 |
| **5** | 84-103 | 5' +G+G+AAG+ACA+GTT+GGG+GAG+A+G+G 3' | 20 | 87 |
| **6** | 104-119 | 5' +C+C+CGG+AGG+GCT+GGG+A+A3' | 16 | 87 |
| **7** | 120-141 | 5'+C+T+GGG+GAA+GCA+AGA+AAG+CC+A+A+G 3' | 22 | 86 |
| **8** | 142-162 | 5'+C+A+AGC+AGC+TGA+ATT+CTT+C+C+G 3' | 20 | 79 |
| **9** | 162-181 | 5' +C+A+GGT+TGG+AGA+TTT+GAA+C+T 3' | 19 | 77 |
| **10** | 182-202 | 5'+G+C+ACA+CTG+CC+TT+AAG+TCC+T+T+T 3' | 21 | 80 |
| **11** | 203-223 | 5' +C+G+GAG+AAA+GGA+GAT+GTCC+T+C+C 3' | 21 | 84 |
| **12** | 220-239 | 5' +C+A+GCT+CTC+TTT+ACT+GGC+G+G+A 3' | 20 | 83 |
| **13** | 240-259 | 5' +A+A+GCA+GCC+TCG+GTC+TTT+G+A+A 3' | 20 | 78 |
| **14** | 260-279 | 5' +A+A+TCC+TCT+CCA+CGT+CAT+C+G+T 3' | 20 | 79 |
| **15** | 280-303 | 5' +T+C+AAA+GCC+TGT+TAA+GGT+TGT+T+G+A 3' | 23 | 80 |
| **16** | 304-328 | 5' +G+C+AGA+TAT+CAC+CGC+TTC+TTG+TC+A+G+T 3' | 25 | 81 |
| **17** | 329-349 | 5' +G+A+G+GAG+CTG+TGC+CAG+AGTG+A+G 3' | 21 | 88 |
| **18** | 350-372 | 5' +C+C+ATT+GTC+CTA+GCA+AGT+GCA+C+T+G 3' | 23 | 84 |
| **19** | 373-394 | 5' +C+C+A+GCA+CCA+AGT+GCT+TTCT+C+A+G 3' | 22 | 84 |
| **20** | 395-411 | 5'+C+G+CCC+CCG+GCA+GAGA+G+G 3' | 17 | 89 |
| **21** | 412-431 | 5' +T+C+TGG+GGT+TTG+GGC+AGT+C+C+T3' | 20 | 88 |
| **22** | 432-451 | 5' +G+C+AAC+TCC+TGC+CTT+AGG+G+T+C 3' | 20 | 80 |
| **23** | 452-473 | 5' +A+A+GAC+CTC+TCC+ATT+GTA+GT+G+C+A 3' | 22 | 78 |
| **24** | 474-496 | 5' +G+A+ACC+TGA+GGT+GCA+GCC+TAA+G+A+C 3' | 23 | 84 |
| **25** | 497-519 | 5' +A+C+GCC+CTT+CCA+GAA+GAT+CTG+A+G+A 3' | 23 | 84 |
| **26** | 520-541 | 5' +G+C+AAC+CAC+AGT+ACC+GCT+GA+A+T+A 3' | 22 | 82 |
| **27** | 542-559 | 5' +A+G+CAG+CTG+CTG+GC+AC+A+G+C 3' | 18 | 88 |
| **28** | 560-580 | 5' +G+G+GGA+GAA+GAG+GTC+TCCA+T+C+A 3' | 21 | 85 |
| **29** | 581-604 | 5' +G+G+CAA+AGA+CGT+TCC+ATT+CAG+AT+G+T 3' | 24 | 81 |
| **30** | 605-631 | 5' +C+T+CAG+TTT+TGA+CAT+TCT+GTA+GACT+C+T+G 3' | 27 | 80 |
| **31** | 632-651 | 5' +T+G+TAG+CTC+TGA+GGT+CGG+A+G+C 3' | 20 | 85 |
| **32** | 652-672 | 5' +G+T+GTG+GAG+TGG+TCC+TTAA+A+G+C 3' | 21 | 78 |
| **33** | 673-691 | 5' +A+G+CAA+CCA+GCT+GCC+TC+T+T+T 3' | 19 | 83 |
| **34** | 682-706 | 5'+G+G+CAG+TT+TC+ATT+AAA+GCA+ACC+A+G+C 3' | 24 | 80 |
| **35** | 692-723 | 5'+A+G+TTC+TTG+AAG+CTT+AAA+GGC+AGT+TTT+CAT+T +A+A 3' | 32 | 77 |
| **36** | 724-749 | 5' +C+T+TAT+AAA+TGG+ATT+CCC+CAA+GGC+C+T+C 3' | 26 | 85 |
| **37** | 750-779 | 5'+A+T+ACA+ACT+TGG+AAA+TGC+ATT+TTC+TAGG+C+T+ C 3' | 30 | 77 |
| **38** | 780-811 | 5'+C+A+GGT+AAA+TTT+TATT+TTG+TAT+CT+CCT+AAG+A+ A+C 3' | 32 | 75 |

| | | |
|---|---|---|
| **Ctrl ASO** | | A*A*C*A*C*G*G*T*C*T*A*T*A*C*G*C |
| **Pnky ASO** | 516-532 | T*A*C*C*G*C*T*G*A*A*T*A*A*C*G*C |

**Table S4. List of primers- genotyping, amplicon primers, RT primers, qPCR primer**

| **Description** | **Sequence (5'-3')** |
|---|---|
| *Pnky* specific RT primer for MaP RT | GCGCAACCACAGTACCGCTGAATA |
| *Pnky* specific RT primer for MaP RT | CAGGTAAATTTTATTTTGTATCTCCTAAGAAC |
| Gene specific RT primer with TruSeq overhang | CAGACGTGTGCTCTTCCGATCTACAATTAATTTTTTCAGGTAAATT |
| TruSeq cDNA ligation | 5'Phos-NNNNNNAGATCGGAAGAGCGTCGTGTAG-3'Bio |
| Amplicon1 Forward primer | GGGAGAAGCAACTTCCTCTG |
| Amplicon 1 Reverse primer | GCGCAACCACAGTACCGCTGAATA |
| Amplicon 2 Forward primer | CCAAACCCCAGAGACCCTA |
| Amplicon 2 Reverse primer | CAGGTAAATTTTATTTTGTATCTCCTAAGAAC |
| *Pnky*_qPCR Forward primer | GGACATCTCCTTTCTCCGCC |
| *Pnky* _qPCR Reverse primer | CACCAAGTGCTTTCTCAGCC |
| GAPDH qPCR Forward primer | GGGAAATTCAACGGCACAGT |
| GAPDH qPCR Reverse primer | AGATGGTGATGGGCTTCCC |
| *Pnky* Genotyping Forward primer | TAAGCTCAAACTCCGGTCCCGGGA |
| *Pnky* Genotyping Reverse1 primer | TCAGGGACAAAGAACCAAAACGAGC |
| *Pnky* Genotyping Reverse2 primer | AATGCTCCCTCTGAGCCTCAATT |
| BAC Genotyping Forward primer | CACCTGCTACCTGATATAGG |
| BAC Genotyping Reverse primer | CCTGCTACCTGATATAGG |
